# Dynamic regulation and requirement for ribosomal RNA transcription during mammalian development

**DOI:** 10.1101/2021.09.22.461379

**Authors:** Karla T. Falcon, Kristin E.N. Watt, Soma Dash, Ruonan Zhao, Daisuke Sakai, Emma L. Moore, Sharien Fitriasari, Melissa Childers, Mihaela E. Sardiu, Selene Swanson, Dai Tsuchiya, Jay Unruh, George Bugarinovic, Lin Li, Rita Shiang, Annita Achilleos, Jill Dixon, Michael J. Dixon, Paul A. Trainor

## Abstract

Ribosomal RNA (rRNA) transcription by RNA Polymerase I (Pol I) is a critical rate-limiting step in ribosome biogenesis, which is essential for cell survival. Despite its global function, disruptions in ribosome biogenesis cause tissue-specific birth defects called ribosomopathies, which frequently affect craniofacial development. Here, we describe a cellular and molecular mechanism underlying the susceptibility of craniofacial development to disruptions in Pol I transcription. We show that Pol I subunits are highly expressed in the neuroepithelium and neural crest cells (NCC), which generate most of the craniofacial skeleton. High expression of Pol I subunits sustains elevated rRNA transcription in NCC progenitors, which supports their high tissue-specific levels of protein translation, but also makes NCC particularly sensitive to rRNA synthesis defects. Consistent with this model, NCC-specific deletion of Pol I subunits *Polr1a*, *Polr1c,* and associated factor *Tcof1* in mice cell-autonomously diminishes rRNA synthesis, which causes an imbalance between rRNA and ribosomal proteins. This leads to increased binding of ribosomal proteins Rpl5 and Rpl11 to Mdm2 and concomitantly diminished binding between Mdm2 and p53. Consequently, p53 protein accumulates, resulting in NCC apoptosis and craniofacial anomalies. Furthermore, compound mutations in Pol I subunits and associated factors specifically exacerbates the craniofacial anomalies characteristic of the ribosomopathies Treacher Collins Syndrome and Acrofacial Dysostosis-Cincinnati Type. Altogether, our novel results demonstrate a dynamic spatiotemporal requirement for rRNA transcription during mammalian cranial NCC development and corresponding tissue-specific threshold sensitivities to disruptions in rRNA transcription in the pathogenesis of congenital craniofacial disorders.

**Significance statement:** RNA Polymerase I (Pol I) mediated rRNA transcription is required for protein synthesis in all tissues for normal growth and survival as well as for proper embryonic development. Interestingly, disruptions in Pol I mediated transcription perturb ribosome biogenesis and lead to tissue-specific birth defects, which commonly affect the head and face. Our novel results show that during mouse development, Pol I mediated rRNA transcription and protein translation is tissue-specifically elevated in neural crest cells, which give rise to bone, cartilage, and ganglia of the head and face. Using new mouse models, we further show that neural crest cells are highly sensitive to disruptions in Pol I and that when rRNA synthesis is genetically downregulated, it specifically results in craniofacial anomalies.

## Introduction

Ribosomal RNA (rRNA) transcription and ribosome biogenesis are critical for cell growth, proliferation, differentiation, and survival. Ribosomes translate cellular proteins and are responsible for the quality and quantity of proteins (1, 2). The ability to modulate translation rates and translation capacity to meet cell-specific needs is regulated in part by the number of ribosomes available to translate mRNAs (2–5). A critical rate-limiting step in ribosome biogenesis is RNA Polymerase (Pol) I-mediated rRNA transcription (6, 7), which accounts for about 60% of all cellular transcription (2, 8) and is integral to increased protein translation during cell growth, proliferation, and other metabolic needs. In mammals, Pol I consists of ten core, one stalk, and two dissociable subunits (9) that transcribes the 47S precursor rRNA, which is then modified, processed, and cleaved into 5.8S, 18S, and 28S rRNAs. These rRNAs, together with 5S rRNA transcribed by Pol III, associate with ribosomal proteins and form the catalytic core of the ribosome (10).

Considering the requirement for Pol I mediated rRNA transcription and ribosome biogenesis in all cell types, it is surprising that defects in these processes result in cancers or tissue-specific developmental disorders known as ribosomopathies (11–14). For example, mutations in the Pol I catalytic subunit *POLR1A* result in Acrofacial Dysostosis Cincinnati Type (AFDCIN) (15), whereas mutations in *POLR1C* and *POLR1D*, shared subunits of Pol I and III, or Pol I associated factor *TCOF1* cause Treacher Collins Syndrome (TCS) (16–18). *TCOF1* encodes the nucleolar phosphoprotein TREACLE, which is involved in rRNA transcription and processing as well as in DNA damage repair (19–21). AFDCIN and TCS present with a range of phenotypes that primarily affect craniofacial skeletal development including micrognathia, cleft palate, and malar hypoplasia (15, 17, 18). The majority of the craniofacial tissues affected in TCS and AFDCIN are derived from neural crest cells (NCC). NCC are a transient progenitor population which arise from the neuroepithelium and then delaminate, proliferate, and migrate into the frontonasal prominences and pharyngeal arches where they differentiate into most of the craniofacial bone and cartilage, among other tissues (22). Therefore, TCS and AFDCIN are considered both ribosomopathies and neurocristopathies due to deficits in ribosome biogenesis and NCC development. Previous work has elucidated the basic functions of the proteins involved in rRNA transcription and ribosome biogenesis in various organisms including yeast and human cell lines (23–25) and demonstrated that mutations in *Tcof1, polr1a, polr1c*, and *polr1d* disrupt NCC development (15, 26–29). However, the mechanisms by which global disruptions in Pol I-mediated transcription result in tissue-specific phenotypes remain poorly understood. In particular, it has not yet been determined 1) why cranioskeletal development is highly susceptible to defects in rRNA transcription and 2) if rRNA transcription is tissue-specifically required during mammalian craniofacial development.

We hypothesized that different cells and tissues have distinct threshold requirements for rRNA transcription, ribosome biogenesis, and protein synthesis to meet their cell-specific needs, and that this leads to distinct cell and tissue-specific threshold sensitivities to deficiencies in rRNA transcription. Given the high incidence of cranioskeletal defects in ribosomopathies, we posited that NCC are one of the cell types that require high levels of rRNA and protein synthesis. We therefore examined the role of Pol I and rRNA transcription in NCC during craniofacial development. Through lineage tracing, expression, and translation analyses, we discovered that neuroepithelial cells and NCC exhibit elevated levels of rRNA transcription which correlate with high levels of protein translation compared to surrounding cells during early embryogenesis.

To understand the intrinsic function of Pol I mediated transcription in NCC, we generated models of Pol I disruption via null and conditional tissue-specific deletion of a catalytic subunit (*Polr1a*), non-catalytic subunits (*Polr1c*, *Polr1d*), and associated factor *(Tcof1*) of Pol I in mice. We discovered that Pol I mediated transcription is essential for cell survival, and that cranial NCC are particularly sensitive to decreased rRNA transcription during early craniofacial development. Pol I subunit and associated factor loss-of-function results in rRNA deficiency, which perturbs the stoichiometric balance between rRNA and ribosomal proteins. This imbalance leads to ribosomal stress, and increased binding of ribosomal proteins, RPL5 (uL18) and RPL11 (uL5), to Murine double minute 2 (Mdm2), a major regulator of p53 activity. Concomitantly, Mdm2 binding to p53 is reduced, which leads to p53 accumulation in the nucleus, and consequently NCC apoptosis and craniofacial anomalies. Thus, global perturbation of rRNA transcription leads to tissue-specific post-translational accumulation of p53 protein, which contributes to the tissue-specificity of developmental ribosomopathy phenotypes. Taken together, our novel work demonstrates the dynamic tissue-specific regulation and requirement for rRNA transcription during craniofacial development that mechanistically accounts for tissue-specific threshold sensitivities to perturbation of rRNA transcription. Finally, our data shows that ubiquitously expressed genes thought to play fundamental housekeeping functions exhibit cell type specific functions, providing novel insights into the roles of rRNA transcription in regulating embryonic development and disease.

## Results

### Cranial NCC have high levels of rRNA and protein synthesis

Mutations in Pol I subunits result in tissue-specific craniofacial anomalies in humans (15, 16, 30). We hypothesized that the underlying cause for these tissue-specific defects is differential transcription of rRNA in NCC during early embryogenesis. We therefore performed ViewRNA® *in situ* hybridization (31) for the 47S pre-rRNA 5’ETS (Fig. 1A), as a measure of nascent rRNA transcription (32, 33) in *Wnt1-Cre; ROSAeYFP* mouse embryos. This transgenic combination lineage labels the dorsal neuroepithelium, including NCC progenitors and their descendants, with YFP (34). At embryonic day (E) 8.5, during NCC formation and early migration, 5’ETS is globally expressed. However, 5’ETS expression was significantly higher in NCC (YFP+ cells) relative to surrounding non-NCC (YFP-cells) (Fig. 1 B, C). At E9.5, during later migration and the onset of differentiation, 5’ETS expression remained higher in NCC versus non-NCC (Fig. 1 D, E), although quantitatively the difference was less than observed at E8.5. This indicates that NCC have endogenously high levels of rRNA transcription at early stages of development while they are in a more progenitor and highly proliferative state compared to surrounding tissues.

**Fig. 1.**
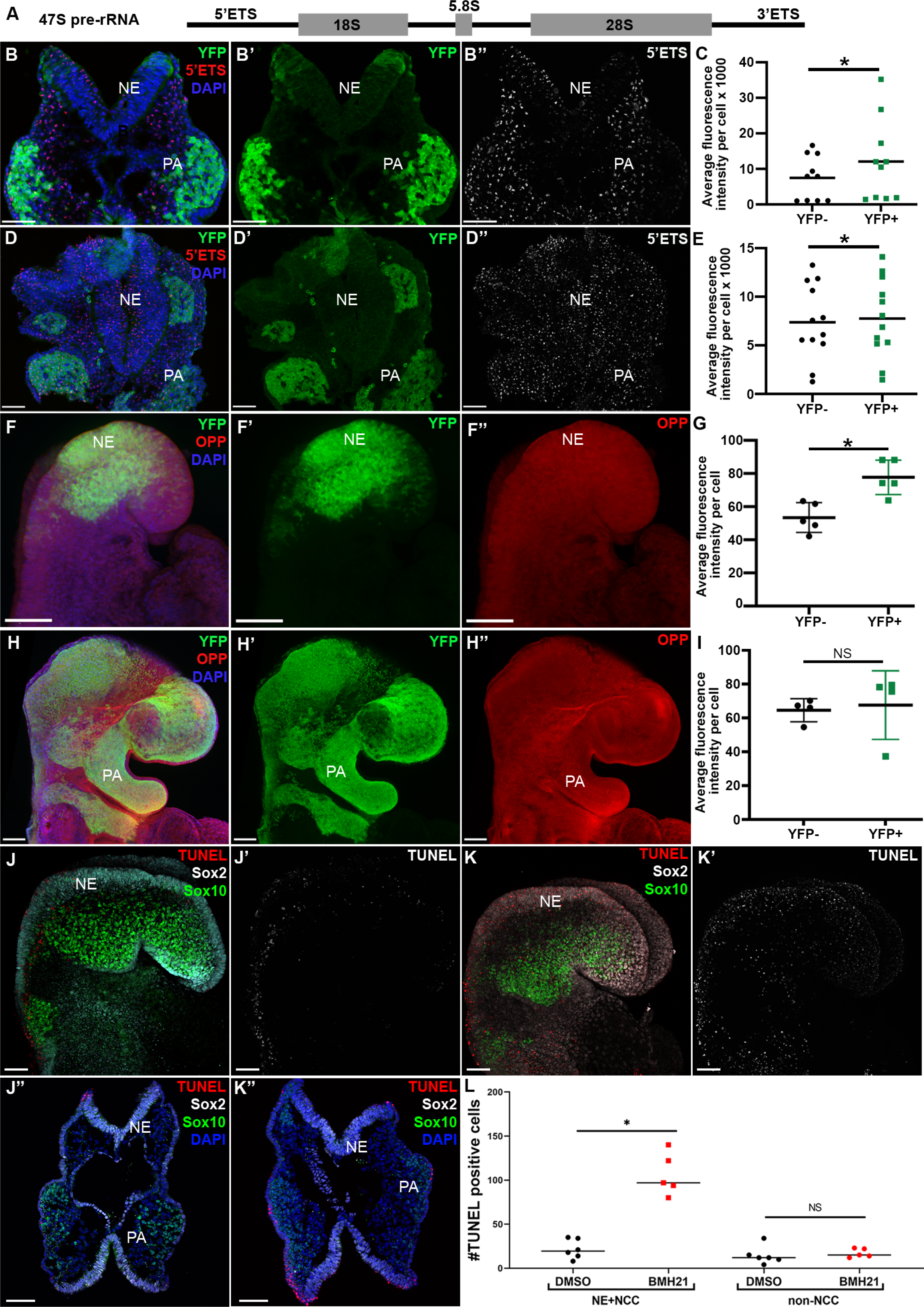
Elevated levels of rRNA synthesis in NCC results in high sensitivity to disruptions in Pol I. (A) Diagram of 47S pre-rRNA showing the 5’ External Transcribed Spacer (ETS), 18S, 5.8S and 28S rRNA components and 3’ETS. (B-E) Fluorescence *in situ* hybridization using ViewRNA^TM^ for the 5’ETS of the 47S pre-rRNA in transverse sections of wild-type *Wnt1-Cre;ROSAeYFP* embryos. At E8.5, 5’ETS expression (red in B, white in B”) is significantly higher in NCC (YFP+; B’) compared to non-NCC (YFP-), quantification in C. At E9.5, 5’ETS expression (D, D”) remains slightly higher in NCC (D’) compared to non-NCC (D’’), quantification in (E). (F-I) Nascent protein synthesis was analyzed via OPP incorporation in wild-type *Wnt1-Cre;ROSAeYFP* embryos. (F) NCC (YFP+) have elevated OPP staining at E8.5 relative to non-NCC (YFP-), quantification in G. (H) OPP staining is comparable between NCC and non-NCC by E9.5, quantification in (I). (J-K”) Disruption of Pol I transcription in wild-type embryos at E8.5 with Pol I inhibitor BMH-21 (K, K’ and K”) results in increased TUNEL positive cells (red in J, J”, K and K” and white in J’ and K’ ) in the neuroepithelium (labeled with Sox2), including the dorsal neuroepithelium where NCC progenitors (labeled with Sox10) are located compared to DMSO treated embryos (J, J’ and J”). (L) Quantification of TUNEL positive cells in DMSO and BMH-21 treated embryos. * indicates p<0.05 using the paired t-test. Abbreviations: NE, neuroepithelium; NS, not significant; PA, pharyngeal arches; NCC, neural crest cells. Scale bar = 100 µm.

Given its importance as a rate-limiting step in ribosome biogenesis, which leads to translation of all cellular protein, we hypothesized that high rRNA transcription in NCC would correlate with elevated protein synthesis. To test this idea, we performed O-propargyl-puromycin (OPP) labeling as a measure of nascent translation (35) in *Wnt1-Cre; ROSAeYFP* embryos, which revealed that cranial NCC have significantly higher protein synthesis compared to other surrounding cells at E8.5 (Fig. 1 F,G), and slightly higher, although not statistically significant, levels of protein synthesis at E9.5 (Fig. 1 H,I). To determine whether increased rDNA transcription and translation correlates with tissue-specific proliferative capacity, we performed 5-bromo-2’-deoxyuridine (BrdU) incorporation in E8.5 wild-type embryos and immunostained transverse sections for BrdU and the mitotic marker phospho-Histone H3 (pHH3). We observed that the neuroepithelium, which includes pre-migratory NCC, is more proliferative than the surrounding mesoderm and endoderm (Fig. S1 A, B). Together, these observations reveal a correlation between high proliferation with elevated rRNA transcription and protein synthesis at E8.5 in neuroepithelial cells and NCC progenitors. Consequently, we posited that the neuroepithelium and NCC, with relatively high levels of rRNA transcription, would be particularly susceptible to cell death upon disruptions in Pol I-mediated rRNA transcription during embryonic development. Culturing E8.5 wild-type mouse embryos for as short as 8 hours with a Pol I inhibitor BMH-21 (36), resulted in apoptosis specifically in neuroepithelial cells and NCC progenitors (Fig. 1 J-L; Fig. S2). Our data therefore demonstrates that endogenously high rRNA transcription and protein translation in the neuroepithelium and NCC progenitors underpins their cell survival-specific threshold sensitivity to disruptions in Pol I.

### *Polr1a, Polr1c, Polr1d,* and *Tcof1* are broadly expressed with elevated levels in the neuroepithelium and pharyngeal arches

To understand if tissue-specific differences in rRNA transcription correlate with differential expression of Pol I subunits during embryogenesis, we examined the expression of *Polr1a, Polr1c, Polr1d* and the associated factor Treacle, which is encoded by *Tcof1,* during early embryogenesis. *Polr1a^+/-^, Polr1c^+/-^,* and *Polr1d^+/-^* mice carrying a gene trap vector with a βGeo cassette in the endogenous locus of each gene were generated (Fig. S3 A), allowing for evaluation of *Polr1a*, *Polr1c*, and *Polr1d* spatiotemporal gene expression by LacZ staining. *Polr1a*, *Polr1c*, and *Polr1d* are broadly expressed at E8.5 and E9.5 (Fig. 2 A-C, I-K) with high levels of expression in the neuroepithelium, and pharyngeal arches, which are the bilateral structures that develop into the jaw and neck (Fig. 2 E-G, M-O). Similarly, Treacle immunostaining of wild-type embryos revealed broad expression in E8.5 and E9.5 embryos, with elevated levels in the neuroepithelium and pharyngeal arches (Fig. 2 D, H, L, P). Furthermore, single-cell RNA sequencing of E8.5 embryos revealed that all the Pol I subunits, including *Polr1a, Polr1c,* and *Polr1d*, as well as *Tcof1* are expressed broadly in progenitor craniofacial cells and tissues (Fig. S4), but each exhibits enriched expression in the neuroepithelium and NCC. Altogether, this suggests that elevated expression of Pol I subunits and associated factor *Tcof1* in the neuroepithelium and NCC contribute to their high levels of rRNA transcription.

**Fig. 2.**
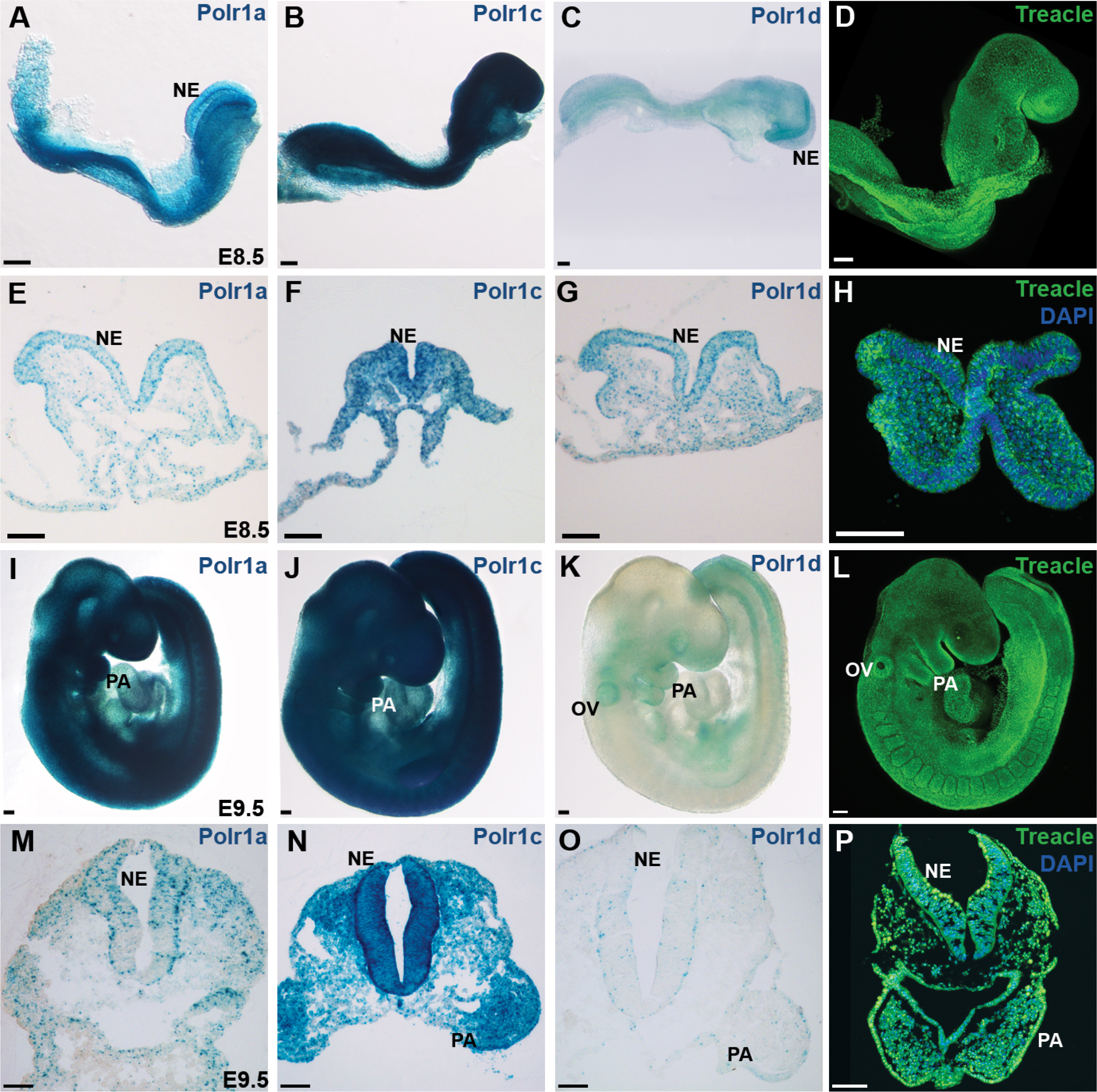
Pol I subunits and associated factor Treacle are broadly expressed during mouse embryogenesis. (A-C) Broad expression of Pol I subunits *Polr1a*, *Polr1c*, and *Polr1d* as observed by LacZ staining in E8.5 embryos. (E-G) Transverse sections through the cranial region indicate high levels of *Polr1a* (E), *Polr1c* (F) and *Polr1d* (G) expression in the neuroepithelium. (I-K) At E9.5, *Polr1a* and *Polr1c* remain broadly expressed (I, J) while *Polr1d* is expressed specifically in the neuroepithelium, pharyngeal arches, otic vesicle and somites (K). Transverse sections through the cranial region at E9.5 indicate higher expression of *Polr1a* (M), *Polr1c* (N) and *Polr1d* (O) in the neuroepithelium and pharyngeal arches compared to surrounding tissues. (D, H, L, P) Immunostaining for Treacle reveals broad expression in whole-embryo and transverse sections of E8.5 and E9.5 embryos, with dynamic elevated levels in the neuroepithelium (H, P). Abbreviations: NE, neuroepithelium; OV, otic vesicle; PA, pharyngeal arches. Scale bar = 100 µm

**Fig. 3.**
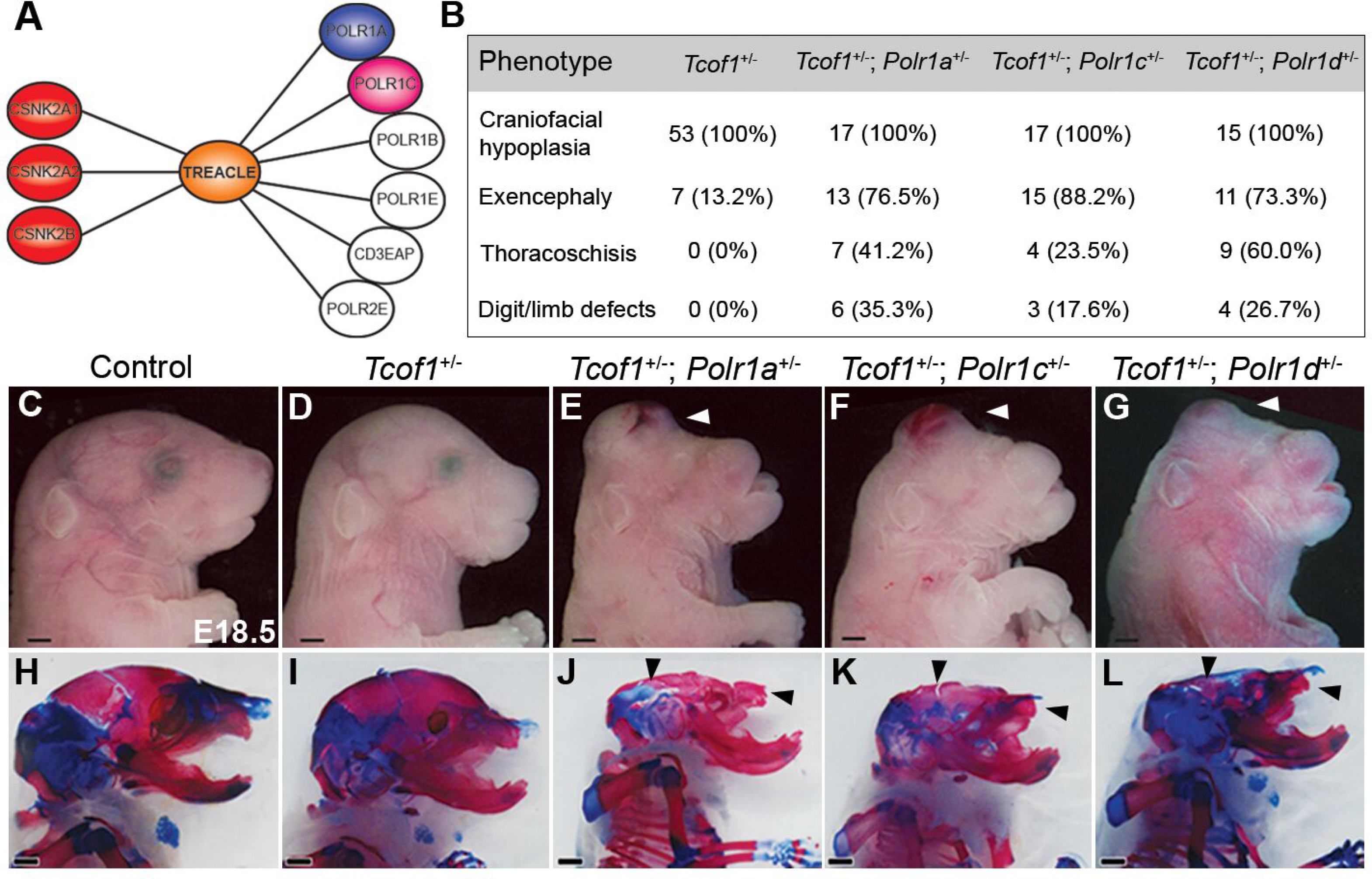
*Tcof1* and Pol I subunits genetically interact, affecting craniofacial development. (A) MudPIT analysis for Treacle-binding proteins recognizes known binding proteins such as Casein kinase II proteins as well as Pol I protein subunits including POLR1A and POLR1C. (B) *Tcof1^+/-^, Tcof1^+/-^; Polr1a^+/-^*, *Tcof1^+/-^; Polr1c^+/-^*, and *Tcof1^+/-^; Polr1d^+/-^* double mutants present with developmental defects with variable penetrance. Table indicates the number of embryos observed with a phenotype. The percentages of the total number of mutants observed are indicated in parentheses. (C-G) Brightfield images of *Tcof1^+/-^; Polr1a^+/-^*, *Tcof1^+/-^; Polr1c^+/-^*, and *Tcof1^+/-^; Polr1d^+/-^* embryos indicate that these double heterozygous mutants exhibit more severe craniofacial defects compared to *Tcof1^+/-^* mutants alone. (H-L) Alizarin red and alcian blue staining for bone and cartilage, respectively, reveals hypoplastic cartilage and/or bone and craniofacial anomalies including smaller maxilla, flattened skulls, and exencephaly in double mutant embryos (J-L) compared to *Tcof1^+/-^* mutants (I) alone. Scale bar = 500 µm.

**Fig. 4.**
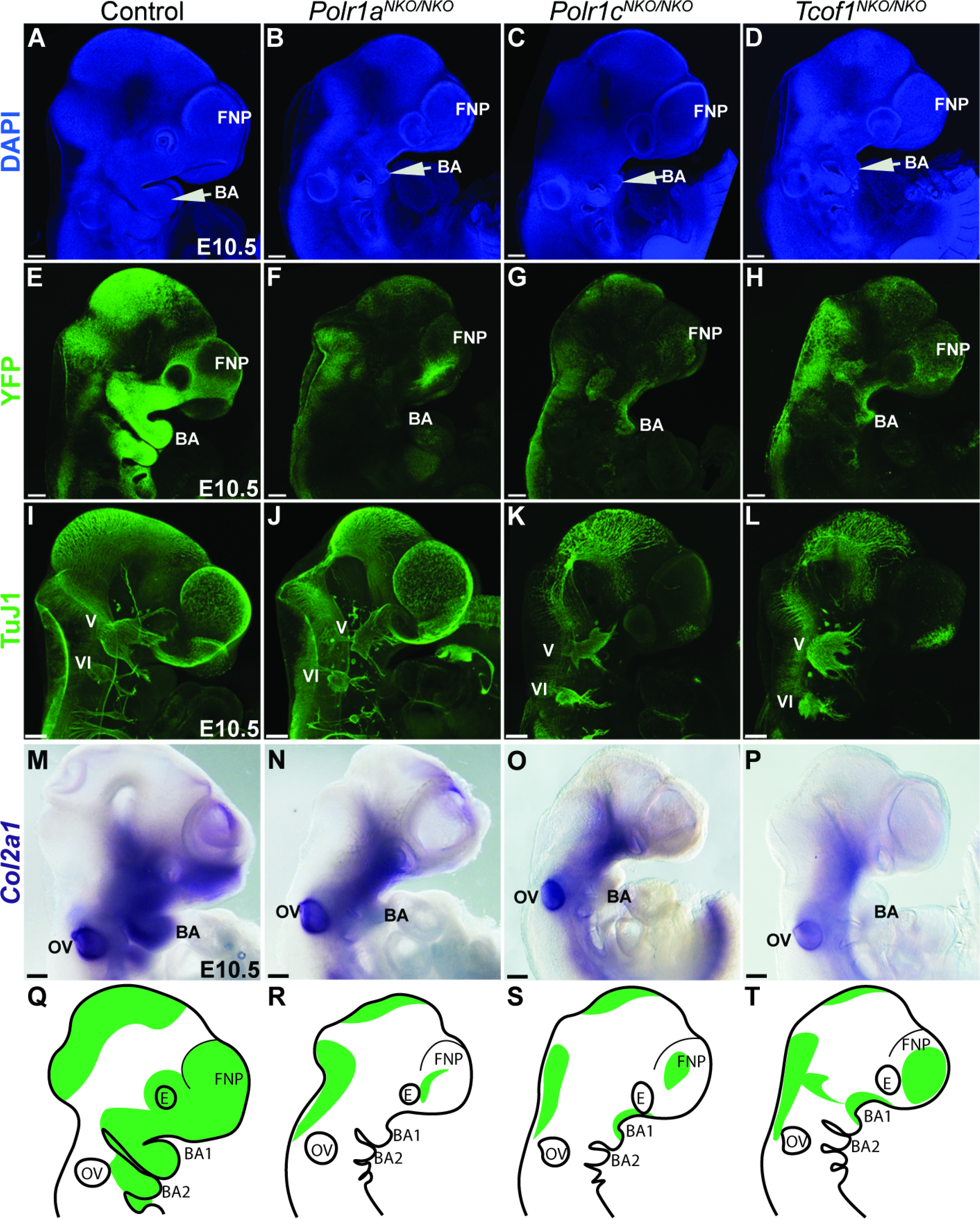
*Polr1a, Polr1c*, and *Tcof1* are required for NCC and craniofacial development in mice. (A-D) DAPI staining at E10.5 shows hypoplastic pharyngeal arches (white arrow) and frontonasal prominences in *Polr1a^NKO/NKO^*, *Polr1c^NKO/NKO^*, and *Tcof1^NKO/NKO^* embryos compared to control embryos. (E-H) *Polr1a^NKO/NKO^*, *Polr1c^NKO/NKO^*, and *Tcof1^NKO/NKO^* were bred into the background of *ROSAeYFP* mice to label the NCC lineage with YFP. YFP staining indicates fewer NCC in the pharyngeal arches and frontonasal prominences in *Polr1a^NKO/NKO^*, *Polr1c^NKO/NKO^*, and *Tcof1^NKO/NKO^* embryos. (I-L) Neuron-specific class III β-tubulin (TuJ1) staining indicates NCC differentiation to neurons and glia is disrupted in *Polr1a^NKO/NKO^*, *Polr1c^NKO/NKO^*, and *Tcof1^NKO/NKO^* embryos. The trigeminal (V) nerve ganglia are hypoplastic in all mutants. (M-P) In situ hybridization for chondrogenesis marker *Col2a1* shows reduced expression especially within the pharyngeal arches in *Polr1a^NKO/NKO^*, *Polr1c^NKO/NKO^*, and *Tcof1^NKO/NKO^* embryos. (Q-T) Schematic figures depicting hypoplastic pharyngeal arches and frontonasal prominences as well as decreased NCC (green) in mutants versus controls. Abbreviations: FNP, Frontonasal prominence; OV, Otic vesicle; PA, pharyngeal arches. Scale bar = 200 µm.

**Fig. 5.**
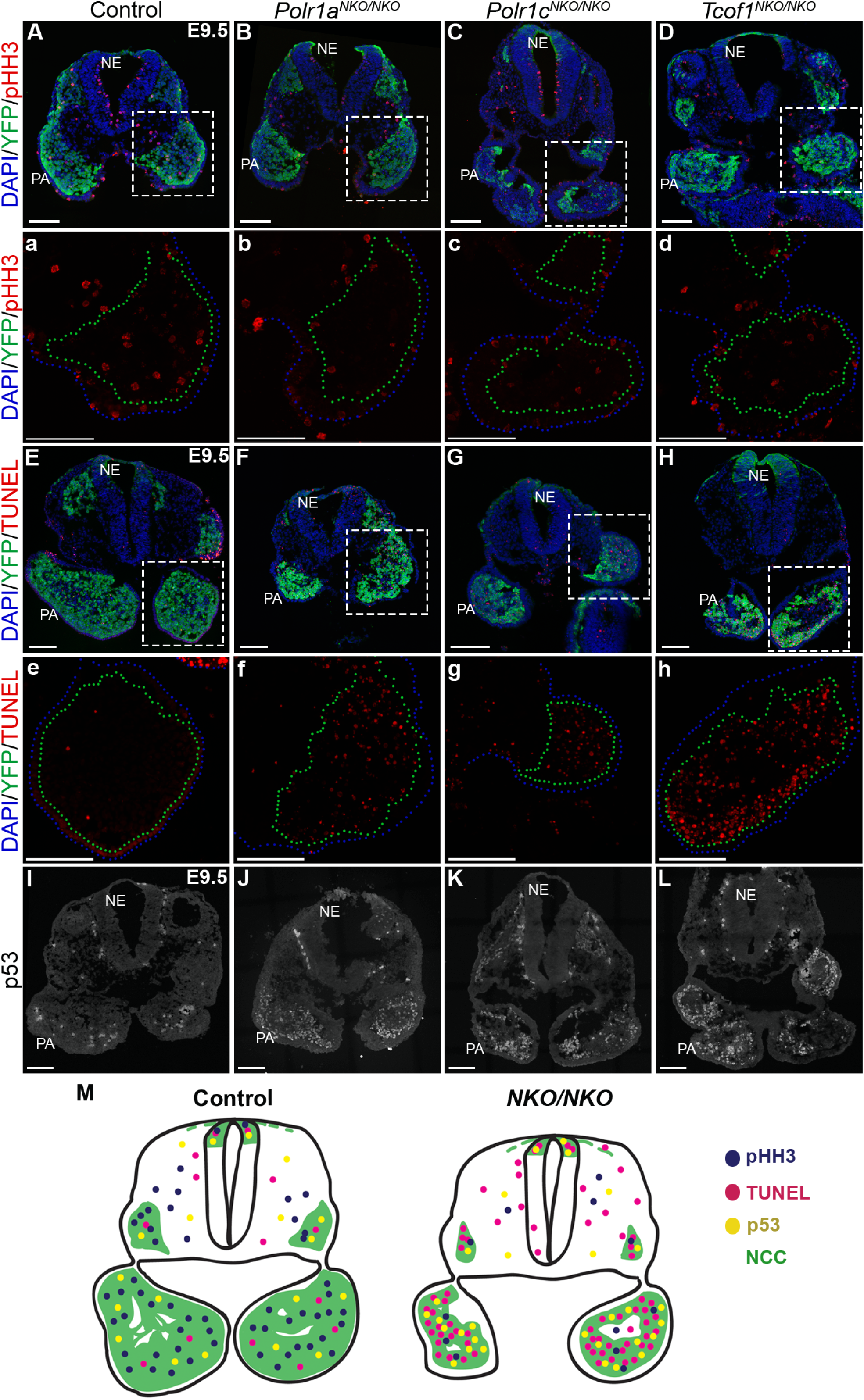
Reduced proliferation and increased p53-dependent cell death underlies the reduced NCC population in *Polr1a^NKO/NKO^*, *Polr1c^NKO/NKO^* and *Tcof1^NKO/NKO^* mice. (A-D) Proliferation (pHH3, red) is reduced in NCC (YFP+) of *Polr1a^NKO/NKO^*, *Polr1c^NKO/NKO^*, and *Tcof1^NKO/NKO^* embryos. (a-d) Higher magnification view of boxed region in A-D. Pharyngeal arches are outlined in green, indicative of the YFP expressing cell boundary and blue, indicative of DAPI labeled cell boundary. (E-H) TUNEL staining shows increased cell death in YFP+ NCC in the pharyngeal arches of *Polr1a^NKO/NKO^*, *Polr1c^NKO/NKO^,* and *Tcof1^NKO/NKO^* embryos at E10.5. (e-h) Higher magnification of the pharyngeal arches, YFP expressing region outlined in green and DAPI with blue. (I-L) Increased p53 staining (white) in the pharyngeal arches in *Polr1a^NKO/NKO^*, *Polr1c^NKO/NKO^*, and *Tcof1^NKO/NKO^* embryos suggests p53-dependent cell death. (M) Summary schematic of control and NCC-specific mutant (representative of *Polr1a^NKO/NKO^*, *Polr1c^NKO/NKO^*, and *Tcof1^NKO/NKO^*) sections depicting decreased levels of pHH3 (blue), increased cell death (pink) and p53 (yellow) levels within NCC (green). Abbreviations. NCC, neural crest cells; NE, neuroepithelium; PA, pharyngeal arches. Scale bar = 100 µm.

To determine the function of individual Pol I subunits during development, heterozygous and homozygous *Polr1a, Polr1c*, and *Polr1d* mutant mice were generated (Fig. S3). *Polr1a^+/-^, Polr1c^+/-^*, and *Polr1d^+/-^* embryos are morphologically indistinguishable from their wild-type littermates at E18.5 and survive to adulthood, indicating that a single copy of each gene is sufficient for proper development in mice (Fig. S3 B). However, *Polr1a^-/-^* (n = 23)*, Polr1c^-/-^* (n=22), and *Polr1d^-/-^* (n=20) embryos are embryonic lethal by E3.5 (Fig. S3 C). Their arrest at the morula stage, and failure to develop into blastocysts and implant demonstrates that these genes are necessary for survival during pre-implantation mammalian development.

### *Polr1a, Polr1c,* and *Polr1d* genetically interact with *Tcof1* during craniofacial development

Given their largely overlapping expression patterns with elevated levels in the neuroepithelium and NCC, together with their shared Pol I associated function in rRNA transcription, we hypothesized that *Polr1a, Polr1c,* and *Polr1d* genetically interact with *Tcof1* during mouse craniofacial development. In support of this idea, we performed Multi-Dimensional Protein Identification Technology (MudPIT) analysis (37, 38) of HEK293T-derived cell lines stably expressing FLAG-tagged TREACLE and found that TREACLE pulled down Pol I subunits, including POLR1A and POLR1C, together with previously known direct targets such as Casein kinase 2 (CSNK2) (Fig. 3A, Fig. S5, Table S1) (39). Thus, POLR1A, POLR1C, and TREACLE interact at a protein level either directly or possibly through a protein-RNA intermediate consistent with being components and associated factors of Pol I.

To functionally test whether these factors interact at a genetic level we generated *Tcof1^+/-^; Polr1a^+/-^, Tcof1^+/-^; Polr1c^+/-^,* and *Tcof1^+/-^; Polr1d^+/-^* double heterozygous mutants. As previously described (26, 40), and compared to controls (Fig. 3 C, H), E18.5, *Tcof1^+/-^* mouse embryos display craniofacial malformations including domed-shaped heads, hypoplasia of the skull, nasal, premaxillary and maxillary bones, together with partially penetrant cleft palate, ear and eye abnormalities (Fig. 3 B, D, I; Fig. S6 A, B), which phenocopies TCS in humans. By comparison, each of the E18.5 double heterozygote mutants exhibit considerably more severe craniofacial defects than found in *Tcof1^+/-^* embryos (Fig. 3 B, D-G).

**Fig. 6.**
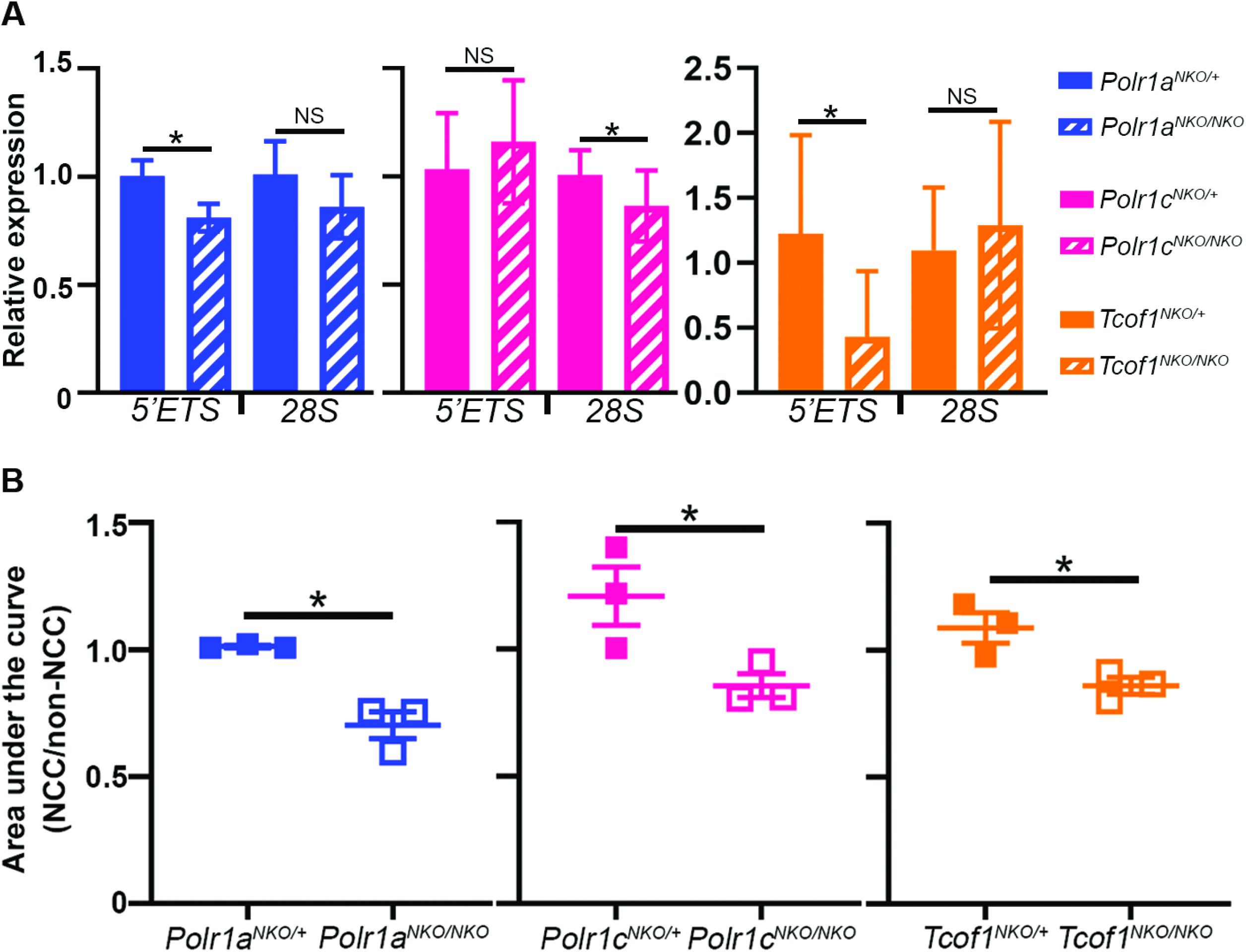
rRNA transcription and total protein are reduced in NCC in *Polr1a^NKO/NKO^*, *Polr1c^NKO/NKO^* and *Tcof1^NKO/NKO^* mice. (A) qPCR for the 5’ETS region of rRNA is significantly reduced in the sorted NCC of *Polr1a^NKO/NKO^* and *Tcof1^NKO/NKO^* embryos, while mature 28S rRNA transcript is not significantly changed. In *Polr1c^NKO/NKO^* embryos, 28S rRNA is reduced while 5’ETS is not significantly affected. (B) Quantification of silver staining demonstrates the total protein in NCC is significantly reduced compared to non-NCC in *Polr1a^NKO/NKO^*, *Polr1c^NKO/NKO^,* and *Tcof1^NKO/NKO^* embryos at E10.5. 2000. YFP+ and YFP-cells of each genotype were used to perform silver staining.

Double heterozygotes display exacerbated craniofacial malformations including fully penetrant cleft palate together with exencephaly and microphthalmia (Fig. 3 B, D-L). Furthermore, alcian blue and alizarin red staining revealed the comparatively more severe hypoplasia and malformation of craniofacial cartilage and bone, particularly of the skull, maxilla and mandible (Fig. 3 I-L; Fig S6 K-W), illustrating the particular sensitivity of craniofacial tissues to perturbations in Pol I function.

Interestingly, the double heterozygous mutant embryos also exhibit variably penetrant developmental anomalies outside of the craniofacial region, which were not observed in *Tcof1^+/-^* embryos. This includes thoracoschisis or omphalocele (fissure of the thoracic or abdominal wall) (Fig. 3B; Fig. S6 A-E), as well as limb and digit anomalies, such as long bone hypoplasia, and abnormal number or short and broad digits (Fig. S6 F-J). While the penetrance of these phenotypes was slightly variable across the double heterozygous mice, we hypothesize that the maternal environment as well as the background of the *Tcof1* mouse strain contributes to some of the phenotypic variability (41). Nonetheless, the exacerbated and complete penetrance of cranioskeletal malformations compared to partial penetrance of other tissue anomalies demonstrates the different threshold sensitivities of distinct tissues to global disruptions in Pol I function. These protein and genetic interactions and the additive effects of their loss-of-function reiterate the importance of tissue specific levels of rRNA transcription and suggest that *Polr1a, Polr1c, Polr1d,* and *Tcof1* function together in rRNA transcription in mammalian NCC during craniofacial development.

### NCC-specific deletion of *Polr1a*, *Polr1c,* and *Tcof1* results in craniofacial defects

Elevated rRNA transcription in NCC progenitors and NCC and the high sensitivity of neuroepithelial and craniofacial tissues to defects in rRNA transcription suggests a cell autonomous role for *Polr1a*, *Polr1c*, and *Tcof1* in Pol I transcription in NCC during early development. We therefore conditionally deleted these factors in NCC during their formation using *Wnt1-Cre* transgenic mice. *Wnt1-Cre* recombinase is expressed in the dorsal neuroepithelium, which includes NCC progenitors beginning at E8.5 (34, 42). We crossed *Wnt1-Cre* mice with *Polr1a^flx/flx^, Polr1c^flx/flx^,* or *Tcof1^flx/flx^* mice to generate NCC-specific knockouts (NKO) of *Polr1a*, *Polr1c*, and *Tcof1* (Fig. S7 A). The levels of *Polr1a, Polr1c,* and *Tcof1* transcripts in NCC were reduced in E9.5 *Polr1a^NKO/NKO^, Polr1c^NKO/NKO^,* and *Tcof1^NKO/NKO^* mutant embryos, respectively, relative to littermate *Polr1a^NKO/+^, Polr1c^NKO/+^,* and *Tcof1^NKO/+^* controls confirming Cre-mediated excision of exons flanked by loxP sites (Fig. S7 B).

**Fig. 7.**
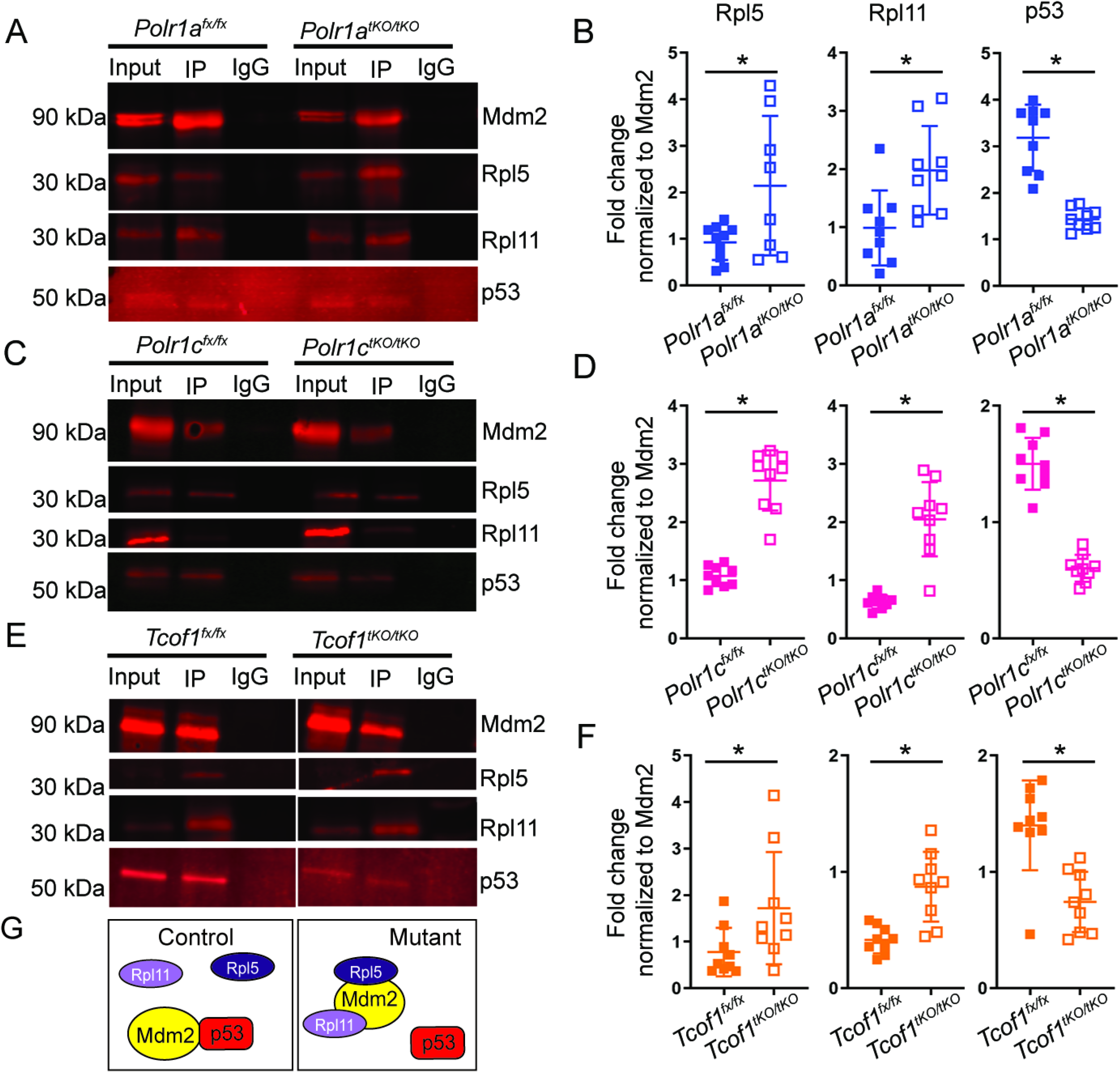
p53 is activated as a result of higher ribosomal protein binding to Mdm2 in mutant mouse embryonic fibroblasts. Mouse embryonic fibroblast cells (MEFs) derived from *Polr1a ^flx/flx^*, *Polr1a^tKO/tKO^* (A-B), *Polr1c ^flx/flx^, Polr1c^tKO/tKO^*. (C-D), *Tcof1^flx/flx^*, and *Tcof1^tKO/tKO^* (E-F) embryos were treated with tamoxifen and used for immunoprecipitation assays. Pull down with Mdm2 and immunoblotting for Rpl5 and Rpl11 revealed increased binding of Mdm2-Rpl5 and Mdm2-Rpl11 in *Polr1a^tKO/tKO^*, *Polr1c^tKO/tKO^*, and *Tcof1^tKO/tKO^* MEFs compared to their respective control MEFs. Conversely, p53 binding to Mdm2 is reduced in *Polr1a^tKO/tKO^*, *Polr1c^tKO/tKO^*, and *Tcof1^tKO/tKO^* MEFs compared to controls, consistent with the increased levels of p53 observed in the *Polr1a^NKO/NKO^, Polr1c^NKO/NKO^*, and *Tcof1^NKO/NKO^* embryos. Band intensities were measured as the ratio between Mdm2 and Rpl5, Rpl11, or p53. (G) Summary schematic showing Mdm2-p53 binding in control and Rpl5, Rpl11-Mdm2 binding in mutant resulting in free p53. * indicates p<0.05, Student’s t-test.

E9.5 *Polr1a^NKO/NKO^, Polr1c^NKO/NKO^,* and *Tcof1^NKO/NKO^* mutants present with visibly hypoplastic frontonasal prominences and pharyngeal arches when compared to littermate controls, a phenotype that worsens considerably by E10.5-E11.5 (Fig. 4 A-D; Fig. S8 A-H). To determine whether this tissue hypoplasia was a consequence of perturbed NCC development, we crossed *ROSAeYFP* into the background of *Polr1a^NKO/NKO^, Polr1c^NKO/NKO^,* and *Tcof1^NKO/NKO^* mice to indelibly label the NCC lineage with YFP (43). This revealed that NCC migrate into the facial prominences and pharyngeal arches in *NKO* mutants by E9.5 (Fig. S8 I-L). However, the smaller facial outgrowths in these mutants appear to correlate with reduced populations of NCC, a phenotype which is even more pronounced at E10.5 in the *NKO* mutants compared to littermate controls (Fig. 4 E-H; Q-T). *Polr1a^NKO/NKO^* embryos have the most severe reduction in the NCC population, consistent with its essential role as part of the catalytic core of Pol I. The NCC population is also severely hypoplastic in *Polr1c^NKO/NKO^* and *Tcof1^NKO/NKO^* embryos as well, although to a slightly lesser degree than *Polr1a^NKO/NKO^* embryos. Reflecting this difference in severity, *Polr1a^NKO/NKO^* embryos die around E11.5, whereas *Polr1c^NKO/NKO^* and *Tcof1^NKO/NKO^* embryos survive until E12.5 and E13.5 respectively (Fig. S7 C, D), conveying the relative importance of *Polr1a*, *Polr1c,* and *Tcof1* in NCC for embryo survival.

NCC differentiate into a wide variety of cell and tissue derivatives including neurons in the peripheral nervous system and osteochondroprogenitors of craniofacial cartilage and bone. To examine NCC differentiation into neurons, we stained for neuron-specific class III β-tubulin (TuJ1) at E10.5. This revealed that *Polr1a^NKO/NKO^* and *Polr1c^NKO/NKO^* mutants exhibit hypoplastic cranial ganglia, especially the trigeminal (V), together with diminished nerve projections compared to littermate controls (Fig. 4 I-K). The trigeminal in *Tcof1^NKO/NKO^* mutants display altered morphology and smaller nerve projections consistent with a reduced population of NCC (Fig. 4 L). The early lethality of *NKO* mutant embryos prevented analysis of NCC differentiation into mature cartilage and bone. Therefore, we investigated the specification of NCC into osteochondroprogenitors. The expression of Sox9, a master regulator of chondrogenesis, and its downstream target, *Col2a1* (44), were both diminished in the facial prominences in E9.5 and E10.5 *NKO* mutants compared to controls (Fig. 4 M-P, Fig. S8 M-T, m-p). The reduced domains of chondrogenic gene expression, especially the first and second pharyngeal arches, and hypoplastic cranial ganglia likely reflect the reduced number of NCC within the arches in *NKO* mutants (Fig. 4 E-H; Fig S8 I-L;). Furthermore, smaller craniofacial prominences and pharyngeal arches (Fig. 4 A-D, Q-T) suggest that *Polr1a, Polr1c,* and *Tcof1* play critical roles in NCC proliferation and/or survival.

### *Polr1a, Polr1c,* and *Tcof1* loss-of-function in NCC leads to increased NCC death

We hypothesized that decreased proliferation and/or increased apoptosis accounts for the reduced NCC population in *NKO* mutants. Transverse sections of E9.5 *Polr1a^NKO/NKO^; ROSAeYFP, Polr1c^NKO/NKO^; ROSAeYFP*, and *Tcof1^NKO/NKO^; ROSAeYFP* embryos were stained for the mitotic proliferation marker pHH3. Quantification revealed that while mutant embryos displayed slightly fewer pHH3+ NCC compared to littermate controls, the differences were not statistically significant at this stage (Fig. 5 A-D; Fig. S9 A). In contrast, TUNEL staining of *Polr1a^NKO/NKO^; ROSAeYFP, Polr1c^NKO/NKO^; ROSAeYFP*, and *Tcof1^NKO/NKO^; ROSAeYFP* mutant embryos revealed increased NCC apoptosis (Fig. 5 E-H; Fig. S9 B), especially within the pharyngeal arches (Fig. 5 E-H). p53 is a well-known mediator of apoptosis (45) and its mRNA level (46) or protein activity (27, 47) has been proposed to underlie tissue-specific defects in the neuroepithelium. We therefore quantified *p53* expression by qPCR and found no significant changes in *p53* transcription between NCC and non-NCC in wildtype embryos (Fig. S10), but *p53* was slightly reduced in the NCC of *Polr1a^NKO/NKO^* mutants compared to *Polr1a^NKO/+^* controls (Fig. S10). This demonstrates that differences in *p53* mRNA levels do not underlie differences in cell death, consistent with previous studies (26, 27). Interestingly, p53 protein is uniformly expressed across different tissues at very low levels in wild-type E8.5 embryos (27), and although *p53* was not affected at the transcript level, p53 protein was tissue-specifically increased in the neuroepithelium and pharyngeal arches in *NKO* mutants compared to their respective littermate controls (Fig. 5 I-L; Fig. S9 C). While p53 protein levels were not significantly increased in *Polr1a^NKO/NKO^* mice at this stage, examination of cell cycle inhibitor and p53 target gene *p21* by qPCR demonstrated a significant increase in *p21* in the NCC of *Polr1a^NKO/NKO^* mutants (Fig. S10 A). This suggests that there may be an effect on proliferation downstream of p53 activation and that the difference in pHH3 observed (Fig. 5 A-B), while not statistically significant, may be biologically significant to the mutant phenotype. To confirm that the overexpression of p53 is biologically relevant in the *Polr1a^NKO/NKO^* mutant mice, we treated these embryos and littermate controls with a p53 inhibitor, pifithrin-α (48). *Polr1a^NKO/NKO^* mice treated with pifithrin-α showed a considerable increase in the volume of the pharyngeal arches in concert with increased YFP+ cells in the arches and frontonasal prominences (n=3/4) compared to *Polr1a^NKO/NKO^* mutants treated with DMSO (Fig. S10 B). This indicates that increased p53 dependent cell death reduces the NCC population in *Polr1a^NKO/NKO^* mutants. However, these pifithrin-α treated *Polr1a^NKO/NKO^* embryos do not survive beyond E12.5, probably because inhibiting p53 does not rescue rRNA synthesis and ribosomal stress. Altogether, our results signify that the NCC population in *NKO* mutants is diminished primarily due to a cell autonomous increase in p53 protein-dependent cell death (Fig. 5 M).

### Excision of *Polr1a, Polr1c,* and *Tcof1* results in decreased rRNA and protein synthesis

Multiple stressors can activate p53 and lead to increased apoptosis or cell cycle arrest and the degree of p53 activation may contribute to the tissue-specificity of developmental syndromes (47). Given the essential role of Pol I subunits and associated factor Treacle in rRNA transcription, we hypothesized that p53 is activated in the *NKO* mutants through a ribosomal stress or nucleolar surveillance response (15, 27, 28). When rRNA transcription is disrupted, this could lead to an imbalance in ribosomal protein to rRNA production, triggering p53 activation. To determine if disruptions in rRNA transcription underlie the increased p53-dependent cell death observed in *NKO* mutants, we analyzed rRNA transcription in fluorescence activated cell (FAC) sorted NCC by quantitative RT-PCR (qPCR). At E9.5, approximately 24 hours after Cre excision, 5’ETS expression was significantly downregulated in *Polr1a^NKO/NKO^; ROSAeYFP* and *Tcof1^NKO/NKO^; ROSAeYFP* NCC when compared to respective control NCC (Fig. 6A). While 5’ETS was not significantly changed in *Polr1c^NKO/NKO^; ROSAeYFP* NCC compared to controls, 28S rRNA, which reflects the level of the precursor 47S transcript and the mature 28S rRNA, was significantly reduced (Fig. 6A). Overall, our data demonstrates that rRNA transcription begins to decrease in *Polr1a*, *Polr1c* and *Tcof1 NKO* mutants as early as E9.5.

Previous studies have shown that reductions in rRNA transcription result in reduced ribosome biogenesis and protein synthesis (29), demonstrating rRNA transcription is a rate-limiting step in ribosome biogenesis. We therefore hypothesized that protein synthesis would be reduced in *Polr1a, Polr1c,* and *Tcof1 NKO* NCC as a consequence of reduced rRNA transcription. Protein was extracted from equal numbers of FAC sorted NCC (YFP+) and non-NCC (YFP-) cells from E10.5 *Polr1a^NKO/NKO^; ROSAeYFP, Polr1c^NKO/NKO^; ROSAeYFP*, and *Tcof1^NKO/NKO^; ROSAeYFP* mutant embryos and their respective controls. Silver staining (49) revealed a significant decrease in total protein in *Polr1a*, *Polr1c,* and *Tcof1 NKO* NCC relative to control NCC (Fig. 6B, Fig. S11). This demonstrates that *Polr1a, Polr1c,* and *Tcof1* loss-of-function in NCC leads to a cell-autonomous reduction in rRNA transcription and total protein, which results in increased p53-dependent NCC apoptosis and consequently craniofacial anomalies.

### *Polr1a, Polr1c* and *Tcof1* deletion results in Rpl5 and Rpl11 binding to Mdm2 and p53 stabilization

To investigate the molecular mechanism by which Pol I disruption activates p53-dependent apoptosis, we generated mouse embryonic fibroblasts (MEFs) from *Polr1a^flx/flx^*, *Polr1c^flx/flx^*, and *Tcof1^flx/flx^* mice crossed to tamoxifen inducible *Cre-ER^T2^*, hereafter referred to as tamoxifen-inducible knockouts (*tKO*). We observed recombination in nearly 70% of the *Polr1a^tKO/tKO^*, *Polr1c^tKO/tKO^*, and *Tcof1^tKO/tKO^* cells, 24 hours after tamoxifen treatment (Fig. S12 D-F). As expected, *Polr1a*, *Polr1c,* and *Tcof1* transcripts were decreased in *Polr1a^tKO/tKO^*, *Polr1c^tKO/tKO^*, and *Tcof1^tKO/tKO^* MEFs compared to *Polr1a^flx/flx^*, *Polr1c^flx/flx^*, and *Tcof1^flx/flx^* control MEFs, 48 hours after tamoxifen induced Cre activation (Fig. S12 A-C).

Consequently, protein synthesis was decreased in the mutant MEFs (Fig. S12 G-I’) while the levels of p53 were increased (Fig. S13), demonstrating the mechanistic equivalency between *tKO* MEFs and *NKO* embryos.

During normal cell growth and proliferation, p53 typically exhibits a short half-life, due in large part to MDM2 (Murine Double Minute 2), which binds to and ubiquitinates p53, targeting it for degradation (50). Mdm2 prevents the accumulation of excess p53 even under conditions of cell stress. However, it has been proposed from *in vitro* studies that when there is an imbalance in the normal stoichiometric ratio of rRNA and ribosomal proteins, free or excess ribosomal proteins, particularly Rpl5 (uL18) and Rpl11 (uL5), bind to Mdm2, inhibiting its function (24, 51–54). rRNA transcription is decreased in *Polr1a*, *Polr1c,* and *Tcof1* NKO embryos; however, western blots showed that the levels of Mdm2, and ribosomal proteins Rpl5 and Rpl11, remain unchanged in *tKO* MEFs compared to controls (Fig. S13). Interestingly, immunoprecipitation followed by immunoblotting revealed increased binding of Rpl5 and Rpl11 to Mdm2, in concert with decreased binding between Mdm2 and p53 in *tKO* MEFs compared to controls (Fig. 7).

These results suggest that disruptions in Pol I-mediated rRNA transcription alter the stoichiometric balance between rRNA and ribosomal proteins, resulting in increased Rpl5 and Rpl11 binding to Mdm2. This diminishes Mdm2 from binding and ubiquitinating p53, leading to tissue-specific p53 accumulation, which can account for the tissue-specific neuroepithelial cell and NCC apoptosis, reduction in NCC, and craniofacial anomalies characteristic of many ribosomopathies (Fig. S14).

## Discussion

rRNA transcription is essential for normal embryo development and our novel mouse knockouts of *Polr1a*, *Polr1c*, *Polr1d* and *Tcof1* demonstrate that Pol I function is critical for pre-implantation whole embryo survival as well as tissue-specific NCC survival. However, why craniofacial development is highly sensitive to perturbations in global rRNA transcription, and Pol I function in humans and animal models (15, 17, 26, 28) remains a critical gap in our knowledge.

Our data demonstrates that rRNA synthesis is tissue-specifically regulated *in-vivo* during mouse embryogenesis and that this correlates with tissue-specific threshold sensitivities to disruptions in rRNA transcription. Quantification of 47S pre-rRNA transcription showed that the neuroepithelium and NCC exhibit endogenously high levels of rRNA transcription compared to surrounding non-NCC (Fig. 1; Fig. S14 A vs C), which is mechanistically underpinned by dynamically enriched expression of *Tcof1*, *Polr1a, Polr1c, Polr1d,* and other Pol I subunit transcripts and protein in the neuroepithelium and NCC in mice (Fig. 2 and S4). This correlates with elevated translation in the neuroepithelium and NCC progenitors, which is necessary to meet high proliferation needs and possibly other demands such as the requirement to translate new proteins for cytoskeletal rearrangement during epithelial to mesenchymal transitions (EMT) (55) (Fig. 1; Fig. S1; Fig. S14A).

Global disruption of Pol I transcription using BMH-21 results in apoptosis specifically in neuroepithelial cells and NCC progenitors in E8.5 mouse embryos (Fig. 1 J-L). Further, craniofacial anomalies are more severe and 100% penetrant in *Tcof1^+/-^; Polr1a^+/-^, Tcof1^+/-^; Polr1c^+/-^,* and *Tcof1^+/-^; Polr1d^+/-^* double heterozygous mutants compared to craniofacial anomalies observed in *Tcof1^+/-^* mutant embryos (Fig. 3). Therefore, taken together with our previous observations in zebrafish (15, 28), this indicates that high rRNA transcription in NCC progenitors leads to their high sensitivity to disruptions in rRNA synthesis while non-NCC derived tissues are affected to a lesser degree (Fig. 3; Fig. S14 B vs D).

Furthermore, Pol I mediated transcription functions in a cell autonomous manner during mouse NCC development. NCC-specific deletion of *Polr1a*, *Polr1c,* and *Tcof1* genes results in NCC autonomous downregulation of rRNA transcription (Fig. 6), leading to increased p53 dependent cell death (Fig. 5) and consequently, craniofacial anomalies (Fig. 4). While mechanistically similar, there are subtle differences between *Polr1a^NKO/NKO^*, *Polr1c^NKO/NKO^*, and *Tcof1^NKO/NKO^* embryos. For example, the NCC population is more severely reduced in *Polr1a^NKO/NKO^* embryos compared to *Polr1c^NKO/NKO^* (Fig. 4), corresponding with previous work in zebrafish (15, 28). *Tcof1^NKO/NKO^* embryos exhibit the least severe phenotype in comparison to *Polr1a^NKO/NKO^* and *Polr1c^NKO/NKO^ .*This is consistent with Polr1a forming part of the catalytic site of Pol I, whereas Polr1c functions to hold Polr1a and Polr1b together but does not form part of the catalytic site, while Treacle is an associated factor that interacts with Pol I (56). Interestingly, while Polr1a is a component of Pol I only, Polr1c is a subunit of both Pol I and Pol III, and therefore may impact Pol III transcription in addition to Pol I, resulting in differences in how rRNA transcription is affected in NCC (Fig. 6A). Modeling of pathogenic variants in *POLR1C* in HeLa cells suggest that the variants associated with TCS primarily affect Pol I function (57), although studies modeling similar pathogenic variants in yeast have indicated that some TCS variants can affect both Pol I and Pol III (25). The roles of Pol III in NCC and craniofacial development is an interesting area for future investigation, and it remains to be determined whether this also involves p53 dependent effects. *Tcof1/*Treacle however, does have additional roles to its function in rRNA transcription, namely in reactive oxygen species-induced DNA damage repair, which when perturbed can also lead to p53 dependent apoptosis (21, 58, 59). Consistent with these functions, antioxidant treatment can ameliorate the craniofacial anomalies in about 30% of *Tcof1^+/-^* mice (21), but a much higher percentage (75%) are rescued with genetic p53 inhibition (27).

Altogether, while there is more to learn about the ribosomal and extra-ribosomal functions of Pol I subunits and associated factors, it remains clear that *Polr1a, Polr1c*, and *Tcof1* are required in NCC and that the phenotypes of the *Polr1a^NKO/NKO^*, *Polr1c^NKO/NKO^*, and *Tcof1^NKO/NKO^* embryos are unified by perturbation of rRNA transcription and an increase in p53-dependent cell death.

While p53 signaling has been implicated in multiple ribosomopathies (45) and developmental syndromes (47), how disruptions in rRNA transcription and ribosome biogenesis result in cell type-specific apoptosis and the molecular mechanism underlying elevated p53 levels in different pathologies remains unclear. (24). Contrary to previous literature that implicates higher transcription of *p53* in the NCC compared to surrounding cells as a potential reason for neurocristopathies (46), we observe that *p53* transcript quantity is similar in NCC and other craniofacial cell types during early embryogenesis (Fig. S10). In addition, previous data shows that p53 protein levels are uniformly low in the neuroepithelium and surrounding tissues in wildtype embryos (27). However, in the absence of Pol I subunits or *Tcof1*, p53 protein is upregulated in a tissue-specific manner (Fig. 5; Fig. S9; Fig. S10). We demonstrate that post-translational p53 activation in *Polr1a, Polr1c,* and *Tcof1* loss-of-function mutants results from an imbalance between rRNA transcription and ribosomal proteins, triggering a nucleolar surveillance response. Excess Rpl5 and Rpl11 bind to Mdm2, limiting Mdm2’s ability to bind to and ubiquitinate p53 (Fig. 7), which lead to p53 protein accumulation and ultimately NCC cell death (Fig. 5; Fig. S14). Further contributing to the tissue specific impact of perturbed rRNA transcription and p53 dependent activation, is that different tissues, including the neuroepithelium and cultured cranial NCC, may be more sensitive to p53 activation (46) and therefore likely to undergo cell death in response to p53 activation (47). Our work therefore suggests that the initial trigger for p53 activation and accumulation in the neuroepithelium in TCS or AFDCIN may be through a nucleolar surveillance mechanism, and that the sensitivity of the neuroepithelium and NCC to p53 activation arises, at least in part, from their elevated requirement for rRNA transcription (Fig. S14). Consistent with this model, the levels of rRNA transcription correlate with their susceptibility to p53-dependent cell death in cancer cell lines. Cancer cells with relatively high levels of rRNA transcription tend to undergo cell death after inhibition of rRNA synthesis whereas cells with relatively low levels of rRNA transcription undergo cell cycle arrest and are more likely to survive.

Altogether, a nucleolar surveillance mechanism may also contribute to other ribosomopathies in which deficiencies in specific ribosomal proteins or increased rRNA transcription (60) are associated with p53-dependent cell death (61, 62). Moreover, it emphasizes the importance of balanced rRNA and ribosomal protein production in the pathogenesis of these pathologies.

Our data suggests that the tissue-specific regulation of rRNA transcription has important implications across multiple diseases and tissue types. While tissue-specific expression and function of ribosomal proteins and pre-ribosomal factors contribute to the pathogenesis of several developmental ribosomopathies (63, 64), the dynamic cell and tissue-specific regulation of rRNA expression during development is not as well understood. Recent studies have observed tissue-specific expression of rRNA in the mouse eye and ovary (32, 33), during forebrain development (65), and during EMT (55, 63). The level of rRNA in these tissues was hypothesized to correlate with levels of proliferation, similar to our data for the neuroepithelium and NCC in E8.5 embryos. Interestingly, the differential levels of rRNA transcription in NCC compared to surrounding cells begins to decrease by E9.5, suggesting that NCC progenitors are more sensitive than NCC at a later developmental stage as they transition from formation, proliferation and migration to differentiation. Consistent with this idea, reductions in rRNA transcription have been observed in association with differentiation in other systems (65–67).

Other factors may also contribute to dynamic tissue-specific rRNA transcription beyond a proliferation and survival versus differentiation demand. This includes epigenetic changes in rDNA (68), rDNA copy number variation (69), tissue-specific expression of variant rRNA alleles (68), or regulation of rRNA synthesis by transcription factors such as Snail1 (55) and Runx2 (70) which are involved in EMT or osteochondroprogenitor differentiation, respectively. Our data suggests that endogenous differential Pol I subunit and *Tcof1* gene expression contributes to the dynamic tissue-specific regulation of rRNA, which underlies the craniofacial defects in TCS and AFDCIN. However, further work is needed to determine the upstream mechanisms that modulate the expression of Pol I and Pol I-mediated transcription, especially in the context of development and disease.

In summary, our novel work has uncovered the dynamic tissue-specific regulation and requirement for rRNA transcription during mammalian embryonic development, which mechanistically accounts for the corresponding tissue-specific threshold sensitivities to disruptions in rRNA transcription, particularly in NCC during craniofacial development. Loss-of-function of Pol I catalytic subunit (Polr1a), non-catalytic subunit (Polr1c), and associated factor (Tcof1) result in similar phenotypes illustrating the conserved mechanisms underpinning the etiology and pathogenesis of Pol I-related craniofacial birth defects in ribosomopathies such as TCS and AFDCIN. Furthermore, we found that the rRNA-Rpl5/Rpl11-Mdm2-p53 molecular pathway which has been previously studied in the context of cancer in yeast and cell lines (24, 53), accounts for the post-translational activation of p53 protein in response to perturbed rRNA transcription. This explains why p53 inhibition is able to suppress apoptosis and rescue craniofacial anomalies in mouse (27) and zebrafish (28, 29) models of rRNA transcription deficiency and raises the possibility that re-establishing the stochiometric ratio between rRNAs and ribosomal proteins could provide a broadly applicable avenue for the therapeutic prevention of ribosomopathies.

## Materials and Methods

### Animal husbandry

All mice were housed in a 16 hour light: 8 hour dark light cycle. All animal experiments were conducted in accordance with Stowers Institute for Medical Research Institutional Animal Care and Use Committee approved protocol (IACUC #2019-097). Transgenic mouse lines were generated at the Stowers Institute for Medical Research Laboratory Animal Facility or by the Virginia Commonwealth University Transgenic/Knockout Mouse Facility (Virginia Commonwealth University IACUC #AM10025). Details of their generation and maintenance can be found in the SI Methods. Mouse Embryonic Fibroblasts were derived from transgenic mice as described previously (71).

### Molecular and phenotypic analysis

Skeletal staining, in situ hybridization, and immunohistochemistry were performed according to previously published methods (72, 73). Description of RNA and proteomic assays along with quantification and statistical analyses are provided in the SI Methods.

## Acknowledgments

The authors thank members of the Trainor lab and Dr Robb Krumlauf for their insights and discussions. We acknowledge Rodney McCay, Lacey Ellington and the Stowers Institute ES Cell and Transgenic Core for generating the *Polr1c^βgeo^* and *Polr1d^βgeo^* mice, and Madelaine Gogol for bioinformatic support. We also thank Mark Miller for illustrating Figure S14. All data needed to evaluate the conclusions in this study are present in the main text, the supplementary materials, and/or the Gene Expression Omnibus (accession no. GSE168351).

## Funding

Stowers Institute for Medical Research (P.A.T)

Kirschstein-NRSA F31 predoctoral fellowship (DE027860) from the National Institute for Dental and Craniofacial Research (K.T.F)

American Association for Anatomy Post-Doctoral Fellowship (A.A.)

Kirschstein-NRSA F31 predoctoral fellowship (DE023017) from the National Institute for Dental and Craniofacial Research (K.E.N.W)

K99 (DE030971) from the National Institute for Dental and Craniofacial Research (K.E.N.W) American Association for Anatomy Post-Doctoral Fellowship (S.D.)

K99 (DE030972) from the National Institute for Dental and Craniofacial Research (S.D.)

NIH National Institute for Dental and Craniofacial Research grant R01DE13172 (L.L. and R.S.)

## Data and materials availability

Original data underlying this manuscript can be accessed from the Stowers Original Data Repository at http://www.stowers.org/research/publications/LIBPB-1604.

## Supplementary Materials

## Supplementary Information

## Materials and Methods

### Mice and animal husbandry

#### Polr1a^+/-^ and Polr1a^flx/flx^

C57BL/6N-*Polr1a^tm1a(EUCOMM)Hmgu/BayMmucd^* mice were obtained from the Mutant Mouse Resource & Research Center and maintained on a C57BL/6 background. The *Polr1a^βgeo/+^* (*Polr1a^+/-^* ) gene trap knockout ready mice were originally generated at the Baylor College of Medicine by injecting ES cell clone HEPD0779_7_B03 into C57BL/6J-Tyr blastocysts. Resulting male chimeras were mated to C57BL/6N females, and the progeny were maintained on a C57BL/6N background. *Polr1a^tm1a^* (*Polr1a^+/-^*) mice were crossed to FlpO (B6.129S4-*Gt(ROSA)26Sor^tm2(FLP*)Sor^*/J, Jax Stock# 012930) mice (1) to generate *Polr1a^flx/+^* mice which were maintained on a C57BL/6 background and then incrossed to generate *Polr1a^flx/flx^* mice (Fig. S2)

### Polr1c^+/-^ and Polr1c^flx/flx^

The *Polr1c^tm1a(KOMP)Wtsi^* ES cells used to generate the *Polr1c ^βgeo/+/-^* (*Polr1c^+/-^*) gene trap knockout ready mouse strain were obtained from the Knock-out mouse project (KOMP) repository. The C57BL/6N parental ES cell line JM8A3.N1 was injected into C57BL/6 blastocysts at the Stowers Institute for Medical Research Laboratory Animal Facility and the *Polr1c^+/-^* gene trap line was established and maintained on a C57BL/6 background. To generate the *Polr1c^flx/flx^* line, mice carrying the FLP recombinase (FLPeR) in the *Rosa26* locus were crossed to the *Polr1c^+/-^* gene trap mice and the resulting *Polr1c^flx/+^* mice were incrossed to homozygosity. The *FLPeR* (B6.129S4-*Gt(ROSA)26Sor^tm1(FLP1)Dym^*/RainJ, Jax stock #009086) mice were obtained from Jackson Laboratory.

### Polr1d ^βgeo/+^ (Polr1d^+/-^)

A C57BL/6 ES cell line (IST10113B8) with a *βgeo* gene trap vector inserted into exon 1 of the *Polr1d* gene was obtained from Texas A&M Institute for Genomic Medicine and injected into C57BL/6 blastocysts at the Stowers Laboratory Animal Facility. *Polr1d^+/-^* mice were maintained on a C57BL/6 background.

### Tcof1^+/-^ and Tcof1^flx/flx^

*Tcof1*^+/-^ mice were generated by insertion of neomycin cassette in exon 1 and maintained as previously described (2) on a DBA background. To generate a conditional allele of *Tcof1,* exon 1 of *Tcof1* was flanked by loxP sites using the targeting vector pTKLNCDL (a gift from Dr. Richard Mortensen) containing a neomycin cassette and loxP sites. The construct was electroporated into ES cells in 129/SvEv ES (HZ2.2) cells and the ES cells that underwent homozygous recombination were cultured and transiently transfected with 7µg of pCMV-cre by electroporation to remove the neomycin cassette, leaving two loxP sites flanking exon 1. This recombination in ES cells was confirmed by PCR and Southern blotting and the cells were injected into C57BL/6 blastocysts to generate *Tcof1^flx/+^* mice by the Virginia Commonwealth University Transgenic/Knockout Mouse Facility (Virginia Commonwealth University IACUC #AM10025). These mice were then backcrossed onto a C57BL/6 background.

### Mef2c-F10N-LacZ

*Mef2c-F10N-LacZ* mice in which LacZ is expressed under the control of a neural crest cell specific enhancer of *Mef2c* were maintained as previously described (3).

### Double Heterozygote Generation

*Tcof1^+/-^* mice were crossed to *Polr1a*, *Polr1c*, or *Polr1d* mice carrying a heterozygous gene trap allele described above.

### Neural Crest Cell knockouts and lineage tracing

*Wnt1-Cre* mice (*H2afv^Tg(Wnt1-cre)11Rth^* Tg(Wnt1-GAL4)11Rth/J, Jax stock #003829) and *RosaeYFP* mice were obtained from the Jackson Laboratory and maintained as previously described (4, 5). *Polr1a^flx/flx^, Polr1c^flx/flx^,* and *Tcof1^flx/flx^* mice were crossed to *Wnt1-Cre* transgenic mice to generate *Polr1a^flx/+^;Wnt1-Cre, Polr1c^flx/+^;Wnt1-Cre,* and *Tcof1^flx/+^;Wnt1-Cre* mice. *Wnt1-Cre* was maintained as a heterozygous allele. To generate *Polr1a^flx/+^;Wnt1-Cre;RosaeYFP, Polr1c^flx/+^;Wnt1-Cre;RosaeYFP,* and *Tcof1^flx/+^;Wnt1-Cre;RosaeYFP* mouse lines used for lineage tracing, *Polr1a^flx/+^;Wnt1-Cre, Polr1c^flx/+^;Wnt1-Cre,* and *Tcof1^flx/+^;Wnt1-Cre* mice were crossed to *RosaeYFP* (B6.129X1-*Gt(ROSA)26Sor ^tm1(EYFP)Cos^/*J, Jax Stock #006148) transgenic mice. *Polr1a^flx/+^;Wnt1-Cre, Polr1c^flx/+^;Wnt1-Cre,* and *Tcof1^flx/+^;Wnt1-Cre* males were crossed to *Polr1a^flx/flx^, Polr1c^flx/flx^,* and *Tcof1^flx/flx^* females, respectively to obtain *Polr1a^flx/flx^;Wnt1-Cre, Porl1c^flx/flx^;Wnt1-Cre,* and *Tcof1^flx/flx^;Wnt1-Cre* embryos, respectively, which are referred to as *Polr1a^NKO/NKO^*, *Polr1c^NKO/NKO^*, and *Tcof1^NKO/NKO^* in this paper.

### Tamoxifen inducible temporal knockouts

Cre-ER^T2^ (B6.129 – Gt(ROSA)26Sor^TM 1(Cre–ERT2)Tyj^/J, Jax stock cat# 008463) mice were crossed to Polr1a^flx/flx^, Polr1c^flx/flx^, and Tcof1^flx/flx^ mice to generate Polr1a^flx/+^;Cre-ER^T2^, Polr1c^flx/+^;Cre-ER^T2^, and Tcof1^flx^/+;Cre-ER^T2^ mice which were subsequently bred to homozygous floxed mice to generate embryos for MEF generation.

The day a vaginal plug was observed in a time mated female was designated as embryonic day (E) 0.5. All mice were housed in a 16 hour light: 8 hour dark light cycle. All animal experiments were conducted in accordance with Stowers Institute for Medical Research Institutional Animal Care and Use Committee approved protocol (IACUC #2019-097).

### Genotyping

To confirm recombination, *Polr1a* and *Tcof1* mice were genotyped according to the primers listed in Supplemental Table 2. Genotyping of all mouse strains was determined using real-time PCR assays with specific Taqman probes designed for each strain (Transnetyx, Inc, Cordova, TN)

### Brightfield imaging

Embryos were imaged on a Leica MZ16 microscope equipped with a Nikon DS-Ri1 camera and NIS Elements imaging software. Manual Z stacks were taken and then assembled using Helicon Focus software. Alterations of brightness and contrast were performed in Adobe Photoshop to improve image clarity and applied equally across the entire image.

### MudPIT

A stable cell line expressing FLAG-tagged TREACLE was generated by transfecting 293-FRT cells with FLAG-Tcof1-pcDNA5/FRT and pOG44 (Invitrogen) using LipofectAMINE 2000(6). Cells were cultured at 37°C in a humidified incubator with 5% CO_2_. Cells were lysed in lysis buffer (50 mM Tris-HCl (pH 7.5), 120 mM NaCl, 0.5% NP-40, 1 mM EDTA, and proteinase inhibitor cocktail (Nacalai tasque)). Following lysis, 2 mM MgC_l2_ and benzonase (50 U/ ml) was added to the whole cell extracts and centrifuged. The supernatant was incubated with agarose beads conjugated with anti-FLAG antibody (Sigma) at 4°C overnight. The following day, the beads were precipitated by centrifugation, washed with lysis buffer, and proteins were eluted with the FLAG peptide (200 μg/ml) in lysis buffer and then precipitated by Trichloroacetic acid. MudPIT was performed to identify interacting proteins as described previously(7, 8).

### Bone and cartilage staining

E18.5 embryos were anesthetized by immersion in ice cold PBS for at least 60 minutes until no reflex movements were observed following a pinch test. The skin and viscera were removed, and the embryos were then fixed in 95-100% ethanol overnight at room temperature or longer at 4°C. Embryos were then stained for bone and cartilage with alizarin red and alcian blue, respectively, as previously described (9). The stained embryos were imaged in 50% glycerol with an MZ16 microscope as described above. Skull measurements were made from the occipital to the nasal bone and mandible measurements were made from the condylar process to the base of the incisor. Measurements of the skull and mandible were taken using ImageJ (NIH, Bethesda, MD).

### BMH-21 treatment

E8.5 embryos were dissected with an intact yolk sac in Tyrode’s buffer and cultured in pre-heated complete media containing 50% DMEM-F12, 50% rat serum, and 1X penicillin/streptomycin in roller culture bottles with 5%CO_2_, 5% O_2_, and 90% N_2_ (10, 11). After 60 minutes of equilibration, 1 µM of BMH-21 (Sigma Aldrich, #SML1183) was added to disrupt Pol I activity. Following 8 hours of incubation, embryos were fixed in 4% PFA/PBS at 4°C overnight. Embryos were stained with Sox2 (1:500, R&D Systems, #AF2018) and Sox10 (1:1000, Abcam, # ab155279) antibodies as well as DAPI following procedures described below. TUNEL assay was performed following manufacturer’s protocol (Roche) described below. The stained embryos were sectioned at 10 µm thickness and imaged using a Zeiss LSM 700 confocal microscope. The sectioned images (three biological replicates and two technical replicates) were analyzed using ImageJ (NIH, Bethesda, MD). Sox2 and Sox10 positive cells were grouped as neuroepithelium and neural crest cells and all other cells were grouped as non-neural crest cells.

### View RNA and in-situ hybridization

E9.5 and 10.5 embryos were harvested in 1X PBS/0.1% DEPC and fixed in 4% PFA in 1X PBS/0.1% DEPC overnight at 4°C. *In situ* hybridization for *Col2a1* (plasmid obtained from Dr. Ralph Marcucio) was performed using standard protocols as previously described (12). Control and mutant embryos were imaged at the same magnification on a Leica stereoscope using a Nikon DS-Ri1 camera. For ViewRNA, embryos were cryosectioned in RNAse free conditions at a thickness of 10μm. Sections were air dried for 20 minutes and washed with 1X PBS. Sections were then dehydrated in 100% ethanol for 5 minutes followed by antigen retrieval with pre-made target retrieval solution (RNAscope® Cat. No. 320850) or citric acid buffer pH 6 (0.1M sodium citrate, 0.1M citric acid in water) at 95°C for 12 minutes. Following proteinase treatment for 5 minutes at room temperature, sections were hybridized and stained per ViewRNA manual instructions (Invitrogen Catalog number: 88-19000).

#### Quantification

Embryo sections were imaged with an LSM-700 upright confocal laser scanning microscope. Prior to intensity quantification, Z stacks of mouse embryo sections stained with antibodies or RNA FISH were sum projected and a uniform background was subtracted based on a manually selected region near the neural tube. Nuclei were detected based on the DAPI signal using a two-dimensional version of the algorithm used for the Click-IT OPP analysis above. Analysis was performed with the aid of the PyImageJ (https://github.com/imagej/pyimagej) interface to Fiji (13) from Jupyter Notebooks (https://jupyter.org/) which are included in supplemental materials. For detection, mask diameters were 20 pixels with a threshold of 20% of the maximum intensity as before. Average intensity measurements were performed using the same mask diameter and two-dimensional histograms and measurements were made of YFP signal vs. ViewRNA signal.

### OPP assay

E8.5 and E9.5 *Wnt1Cre;YFP* embryos were cultured in 1:1000 OPP in DMEM-F12 culture media for 1 hour, followed by the Click-IT reaction per Click-iT^TM^ Plus OPP Protein Synthesis Assay Kit (Invitrogen, Catalog #C10457) manual instructions. MEFs treated with tamoxifen and DMSO were treated with 1:500 OPP for 3 hours, followed by the Click-IT reaction.

#### Quantification

Embryos were imaged with an LSM-700 upright confocal laser scanning microscope. OPP intensity levels were quantified using custom ImageJ (NIH, Bethesda, MD) plugins (13). Detection of nuclear positions from the DAPI signal was performed using a maximum mask approach. This method detects the maximum intensity in the 3D image and then masks out a spheroidal region around it with an XY diameter of 25 pixels and z diameter of 15 slices. The maximum intensity is then found again and masked again repeatedly until there are no maximum pixels above a specified threshold. The threshold was set as 20% of the maximum DAPI intensity in the image. This algorithm does not find the nuclear positions perfectly, but it does provide a measurement proportional to the nuclear density. Average intensities in the YFP and OPP channels were then measured centered at the nuclear positions in a spheroid with an XY diameter of 20 pixels and a z diameter of 10 slices. This smaller measurement region ensures that slightly overlapping nuclei do not significantly influence the measurement. Two dimensional logarithmically binned histograms were made of the GFP signal (denoting the neural crest population) vs. the OPP signal. Those histograms showed clear positive and negative GFP populations allowing for manual drawing of rectangular gates for these populations followed by simple average intensity per cell calculations. In some cases, laser powers were adjusted during the signal acquisition and those values were corrected for in the measurement of the intensities.

### β-galactosidase staining

E8.0-E9.5 embryos were collected and fixed in 2% PFA/0.2% glutaraldehyde in PBS for the following time durations: E8.5-E9.0 for 15 minutes and E9.5 for 30-45 minutes at 4°C. Embryos were rinsed with PBS and stained according to manufacturer’s protocol (Millipore #BG-6-B, #BG-7-B, #BG-8-C). Embryos were then fixed again in 4%PFA/PBS at 4°C rocking overnight and washed in PBS for whole embryo brightfield imaging. For sections, embryos were rinsed and immersed into 30% sucrose/PBS overnight at 4°C. The following day they were submerged into 1:1 30% sucrose/OCT and then embedded in OCT and cryosectioned at a thickness of 10μm. Sections were then imaged on an Axioplan 206 Std microscope with Micro-manager 1.4, Win 10 software.

### Immunostaining

Embryos were harvested at the desired developmental stages in 1X PBS and fixed in 4% PFA/1X PBS at 4°C overnight with the exception of p53 staining which required fixation in 4% PFA/1X PBS at 4°C for 3 hours. For whole embryo staining, the embryos were dehydrated through an ascending methanol series into 100% methanol and stored in -20°C overnight. Next, embryos were treated with 4:1:1 Methanol: DMSO: Hydrogen Peroxide and rehydrated through a descending methanol series into PBS. Embryos were blocked with 2%BSA/2% goat serum prior to staining. For section-staining, the fixed embryos were cryosectioned transversely at 10 µm thickness, followed by blocking solution and staining as previously mentioned. For p53 staining, antigen retrieval was performed by immersing the sections in pre-warmed citric acid buffer (pH6, 0.1M sodium citrate, 0.1M citric acid in water) and incubated at 80-90°C for 30 minutes. Sections were then permeabilized with 0.5% TritonX-100 in PBS followed by 3% BSA blocking solution. Primary antibodies used were: Tcof1 (1:1000, Abcam# ab65212), Sox9 (1:200, Abcam # ab185966), TuJ1 (1:500, Covance Research products, # MMS-435P), GFP (1:500, Life Technologies #A6455), phospho-histone H3 (1:2000, Millipore # 06-570) and p53 (1:100, Cell Signaling Technology #2524S). The embryos and sections were counter-stained with DAPI (Sigma-Aldrich #D9564) to visualize the nuclei. Embryos were imaged with an LSM-700 upright confocal laser scanning microscope. Confocal optical slices were collected and maximum-intensity projections of stacks were made with Zeiss LSM software.

#### Quantification

pHH3 measurements were performed similarly to the ViewRNA measurements. The DAPI channel was Gaussian blurred with a standard deviation of 5 pixels and the diameter for nuclear detection was 45 pixels with a threshold at 15% of the maximum intensity. The measurement diameter was 30 pixels. To determine the fraction of positive cells, the intensity of the brightest positive cells was measured as the average intensity of the four brightest cells in the image. This method assumes that there are at least 4 positive cells in each image which we have confirmed by visual inspection. The cutoff for positive cells was then set at 20% of that positive value and the fraction of cells above that cutoff was measured. Measurements are only reported for the YFP positive population. p53 measurements were performed identically to pHH3 but including all cells in the section. For statistical analysis, fluorescence intensities of *Polr1a^NKO/NKO^, Polr1c^NKO/NKO^,* and *Tcof1^NKO/NKO^* were compared to *Polr1a^NKO/+^, Polr1c^NKO/+^,* and *Tcof1^NKO/+^*, respectively.

### TUNEL staining

Following overnight 4%PFA/PBS fixation, embryos were washed in 1X PBS and then dehydrated through an ascending methanol series into 100% methanol and stored at -20°C overnight for wholemount staining. Embryos were then rehydrated through a descending methanol series into PBS. Alternatively, after fixation, embryos were placed into 30% sucrose/PBS overnight at 4°C and then embedded in OCT for cryosectioning. Embryos and cryosections were permeabilized in 0.1% sodium citrate/PBT (0.1% TritonX in 1X PBS) for 10 minutes at room temperature. Samples were then washed in PBS and were then incubated with 1:19 TUNEL enzyme: buffer (Roche) at 37°C in the dark for 1-2 hours and then counter-stained with DAPI (Sigma-Aldrich #D9564). Embryos were imaged with an LSM-700 upright confocal laser scanning microscope similar to the immunostaining.

#### Quantification

TUNEL measurements were performed similarly to the ViewRNA measurements. The fraction of positive cells was measured as the fraction of cells with an intensity above 10,000 units, a level corresponding approximately to the level of positive cells seen in the image. Measurements are reported only for the YFP+ population.

#### BrdU labelling

To analyze cell proliferation, E8.5 pregnant mice were injected intraperitoneally with BrdU at 0.1mg/kg of body weight. After 30 minutes of incubation, mice were sacrificed. For detection of BrdU-positive cells, transverse cryosections were incubated with 1M HCl for 30 minutes at 37°C after immunostaining with pHH3 (Millipore # 06-570, dilution 1:500), and following secondary antibody incubation. BrdU-positive cells were detected by immunostaining using a rat anti-BrdU antibody (Abcam, dilution 1:200). The number of BrdU+ and pHH3+ cells in neuroepithelium, mesoderm and endoderm were counted (three biological replicates, five technical replicates). Fluorescence microscopy was performed on a LSM5 PASCAL confocal microscope (Carl Zeiss).

### Cell sorting

E9.5 and E10.5 embryos were dissected in Tyrode’s buffer and yolk sacs were saved for genotyping. Control and mutant embryos positive for YFP were used for the cell sorting. The embryos were incubated at 37°C for 5 minutes with TypLE (Gibco) and vortexed for 10 seconds. This cycle of incubation and vortexing was repeated for a period of 15-20 minutes to obtain a single cell suspension, following which TypLE was quenched with fetal bovine serum. The cells were then centrifuged at 200 rcf for 10 minutes. The supernatant was discarded, and the cells were resuspended in PBS. 1µl of 100 µg/ml propidium iodide was added to gate viable cells and the cells were sorted using a FACSMelody (BD Biosciences). YFP+ and YFP-live cells were immediately processed for RNA and protein isolation (2000 cells each). Downstream analysis was performed after confirmation of genotypes.

### RNA isolation, cDNA preparation and qPCR

RNA was extracted from sorted YFP+ and YFP-cells from control and mutant embryos using the Qiagen miRNeasy Micro Kit. RNA was tested for quality on the Agilent 2100 Bioanalyzer and only RNA samples with a RIN score greater than 8.0 were used. The Superscript III Kit (Invitrogen) was used to synthesize cDNA for qPCR using random hexamer primers. qPCR was performed on ABI7000 (Thermo QuantStudio 7) using Perfecta Sybr Green (Quantbio # 95072-250). Primers are listed in Supplemental Table 2. Primers for *Polr1a* were designed on exons upstream of the floxed exon while primers for *Polr1c* and *Tcof1* were designed on the floxed exon. No template controls were run as negative controls. ΔΔCt method was used to calculate fold change. Student’s t-test and ANOVA were used for statistical analysis and significance was determined based on p < 0.05.

### Silver staining

2000 YFP+ cells and YFP-cells from controls and mutants were sorted using a FACSMelody (BD Biosciences) sorter as mentioned previously. Cells were lysed at 4°C for 30 minutes using 20 µl of lysis buffer containing Tris pH 8.0, sodium chloride, sodium deoxycholate, SDS, NP-40, and protease inhibitor. Following lysis, the cells were centrifuged at 13,000 rpm at 4°C for 30 minutes. 1X Laemmli buffer (loading buffer) was added to the extracted protein and denatured at 95°C for 5 minutes. The protein was then loaded onto 4-20% gradient SDS-PAGE gels and run in an electrophoresis unit for 90 minutes at 90V. The gel was then stained using a Pierce Silver Stain kit (ThermoFisher Scientific, #24612) following the manufacturer’s instructions. Band intensities were measured as area under the curve using ImageJ.

### Mouse embryonic fibroblast derivation

Mouse embryonic fibroblast cells (MEFs) were derived from E13.5 and E14.5 *Polr1a^flx/flx^, Polr1a^flx/flx^;Cre-ER^T2^*, *Polr1c^flx/flx^*, *Polr1c^flx/flx^;Cre-ER^T2^, Tcof1^flx/flx^* and *Tcof1^flx/flx^;Cre-ER^T2^* embryos as described previously (14). Cells were cultured in a complete media containing DMEM, 30% FBS, 1X L-Glutamine, 1X Non-essential amino acids, and 1X 2-mercaptoethanol and kept in passage for 3-5 generations. For deletion of *Polr1a*, *Polr1a^flx/flx^;Cre-ER^T2^* MEFs were treated with 5 µM tamoxifen dissolved in DMSO, while *Polr1c* and *Tcof1* deletion was performed by treating *Polr1c^flx/flx^;Cre-ER^T2^* and *Tcof1^flx/flx^;Cre-ER^T2^* MEFs with 1µM tamoxifen. *Polr1a^flx/flx^*, *Polr1c^flx/flx^* and *Tcof1^flx/flx^* MEFs treated with tamoxifen were used as controls. The treatment was performed for 24 hours and the cells were allowed to recover for 24 hours. All experiments were performed 48 hours post tamoxifen induction with three biological replicates of MEFs derived from three mutants as well as in three technical replicates. RNA isolation and qPCR were performed using the same approach and primers as above.

### Western blot

MEFs treated with tamoxifen were lysed using lysis buffer and western blot was performed using standard protocols as described previously (15). Protein quantity was estimated via a BCA assay. Antibodies used were p53 (1:500, Cell Signaling Technology, #2524S), Rpl5 (1:1000, Cell Signaling Technology, #51345), Rpl11 (1:1000, Cell Signaling Technology, #18163), Mdm2 (1:500, Cell Signaling Technology, #86934) and ɣ-Tubulin (1:1500, Millipore Sigma, #T6557). Western blots were imaged and quantified using a CLx-Scanner (Li-COR) and Odyssey Software. For quantification, band intensities for Rpl5, Rpl11, Mdm2 and p53 were compared to housekeeping control ɣ-Tubulin. Student’s t-test was performed for statistical analysis.

### Immunoprecipitation

MEFs were cultured on a T75 plate and harvested following tamoxifen treatment. Immunoprecipitation was performed as previously described (15). Briefly, the cells were homogenized in 500 µl lysis buffer containing Tris pH 8.0, Sodium Chloride, SDS, Sodium deoxycholate, NP-40 and protease inhibitor. The homogenized mixture was then incubated with overhead rotation for 30 minutes at 4°C followed by centrifugation at 13,000 rpm at 4°C. The lysate was then divided into two tubes with of equal protein content, one for incubation with Normal Rabbit IgG and the other for incubation with Mdm2 antibody (Cell Signaling Technology, #86934). 2 µg of antibody was used per mg of protein for immunoprecipitation. 10% volume of the lysate used for immunoprecipitation was collected separately to be used for the control input lane for western blot analysis. The lysate-antibody mix was incubated at 4°C overnight with overhead rotation with a speed of 40 rpm. The following day pre-washed Dynabeads were incubated with the lysate-antibody mix at 4°C for 4 hours. The beads were then washed and eluted with 2X Laemmli buffer at 95°C. The eluted protein was then used for SDS-PAGE and western blot. The protein bands were detected using antibodies against p53 (Cell Signaling Technology, #2524S), Rpl5 (Cell Signaling Technology, #51345), Rpl11 (Cell Signaling Technology, #18163) and Mdm2 (Cell Signaling Technology, #86934). For each biological replicate, cells from one T75 culture plate were used for immunoprecipitation. The experiment was performed in three biological and three technical replicates. For quantification, band intensity of Rpl5, Rpl11, and p53 were compared to Mdm2 in both control and mutant cells. Student’s t-test was performed for statistical analysis.

### Drug Treatment

Pregnant dams were injected intraperitoneally for four consecutive days from E6.5-E9.5 with 3 mg of pifithrin-α per kg of body weight of the mouse. For control experiments, pregnant mice were injected with 200 ul of 50% DMSO for the same period of time. At E10.5, the embryos were dissected and immunostained for GFP.

### Single cell RNA sequencing

#### Tissue collection

6 *Mef2c-F10N-LacZ* (3) and 6 *Wnt1-Cre;RosaeYFP* (5, 16, 17) mice were collected at E8.5. Cranial tissues were manually dissected and incubated in 0.25% Trypsin+EDTA in a 37°C water bath for 1 minute and dissociated through gentle repetitive pipetting about 10 times. The tube was then incubated again in a 37°C water bath for another minute prior to the addition of cold FBS to block further reaction with Trypsin activity. Samples were then centrifuged at 1600 rcf at 4°C for 15 minutes. The supernatant was discarded and the cells were resuspended in 200 µl PBS+2% FBS. Cells were centrifuged again and resuspended in 40 µl PBS. Genotyping was performed after cell dissociation which indicated that 3 out of the 6 *Mef2c-F10N-LacZ* embryos were LacZ positive and that 4 out of the 6 *Wnt1-Cre;RosaeYFP* embryos were YFP positive.

#### Processing and Sequencing

Dissociated cells were assessed for concentration and viability using a Luna-FL cell counter (Logos Biosystems). The cells were confirmed to have at least 70% viability and 12,000-15,000 cells per sample were loaded on a Chromium Single Cell Controller (10x Genomics). Libraries were prepared using the Chromium Next GEM Single Cell 3’ Library & Gel Bead Kit v3.1 (10x Genomics) according to manufacturer’s directions. Resulting short fragment libraries were checked for quality and quantity using a Bioanalyzer (Agilent) and Qubit Fluorometer (ThermoFisher). Libraries were pooled at equal molar concentrations and sequenced on an Illumina NovaSeq 6000 S1 flow cell with the following paired read lengths: 28 bp Read1, 8 bp I7 Index and 98 bp Read2.

#### Data processing

Raw sequencing data was processed using Cell Ranger (v3.0.0, 10x Genomics) to generate gene-level counts for each cell in each sample. Genes with counts in less than three cells were removed from the dataset. Mitochondrial percentages and feature count distribution were used as criteria for cell quality control. The percent mitochondria threshold was set to keep 75% of cells of the *Mef2c-F10N-LacZ* sample (mito ≤ 10.93%). The same threshold was applied to the *Wnt1-Cre;RosaeYFP* sample, keeping 86% of the cells. In addition, cells with feature counts of > 10,000 or < 500 were also excluded from the analysis. The final dataset used for analysis consisted of 21,190 cells (12,498 cells for *Wnt1-Cre;RosaeYFP* and 8,692 for *Mef2c-F10N-LacZ*) and 29,041 genes and is available at the Gene Expression Omnibus (accession no. GSE168351).

The Seurat package (v3.1.1)(18) was used to normalize data via the SCTransform method (19). Mitochondrial percentage was regressed out during normalization. For clustering, 3000 highly variable genes were selected, and the first 46 principal components based on those genes were used to identify 7 clusters at a resolution of 0.05 using the shared nearest neighbor method. The identities of clusters were determined by the differential gene expression of classic markers for each tissue type. Neuroepithelial cells were identified by high expression of classic markers, *Sox2* and *Sox1*. Neural crest cell reporters, *LacZ* and *pEYFP*, and *Sox10* were used to cluster NCC. Embryonic blood cells were clustered based on *Hba-*x expression and endothelial cells based on *Kdr* expression. *Cdh1* expression was used to identify non-neural ectoderm and *Tbx1* expression was used to cluster endodermal and mesodermal cells. Data was visualized in reduced dimensionality using UMAP.

The expression value for each gene was standardized by subtracting the gene’s mean expression and dividing by its standard deviation. For instance, a value of -1 would imply that the value is one standard deviation below the mean expression for that specific gene.

**Fig. S1.**
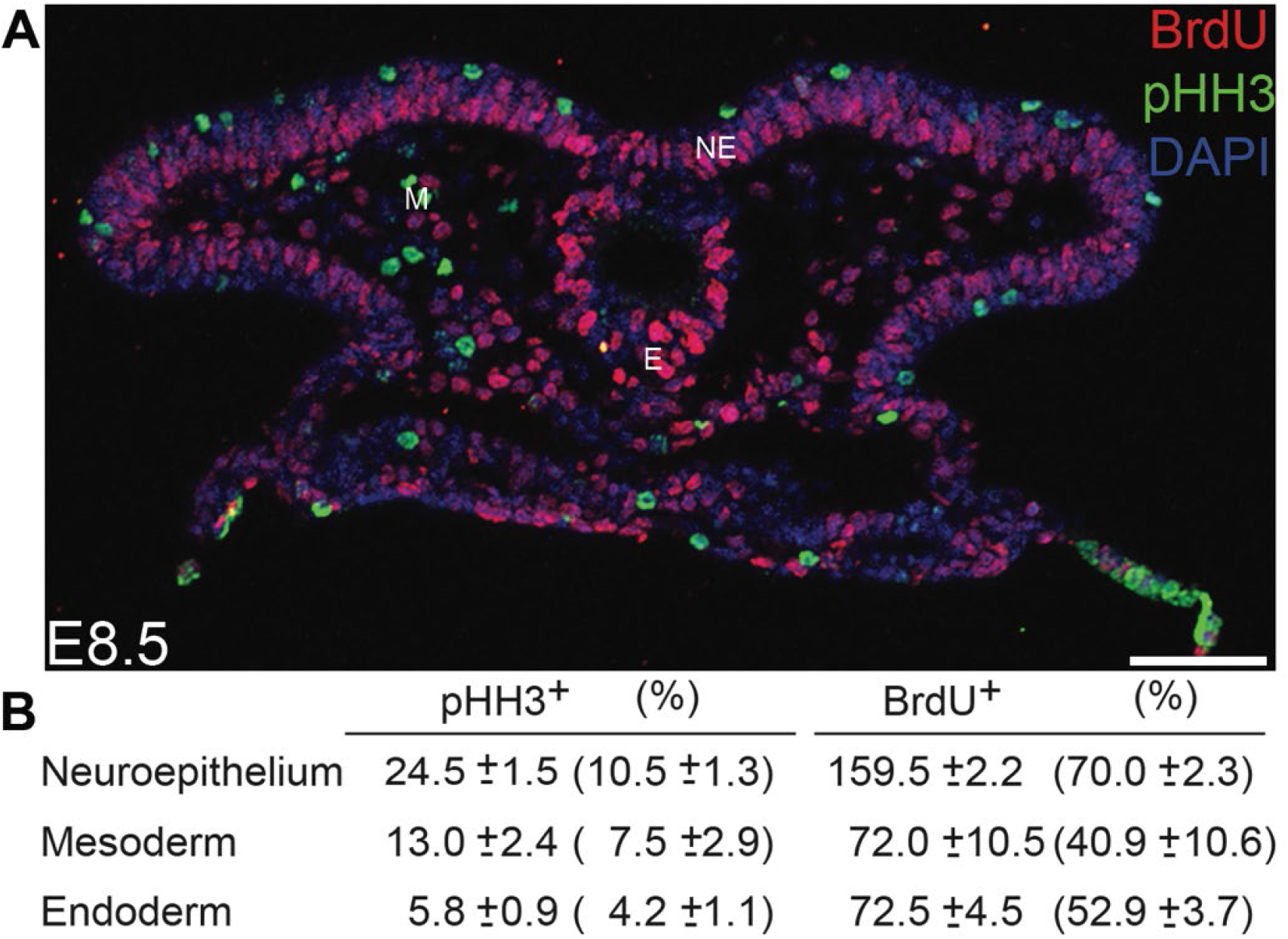
The neuroepithelium is highly proliferative. (A) A higher number of cells in the neuroepithelium, which includes premigratory NCC, are pHH3 and BrdU positive compared to surrounding mesoderm and endoderm cells, indicating that the neuroepithelium is more highly proliferative at E8.5 in wild-type embryos. (B) Quantification of pHH3 and BrdU positive cells in E8.5 craniofacial tissue from three biological replicates and five technical replicates. Scale bar = 80 µm. Abbreviations: NE, neuroepithelium; M, Mesoderm; E, Ectoderm.

**Fig. S2.**
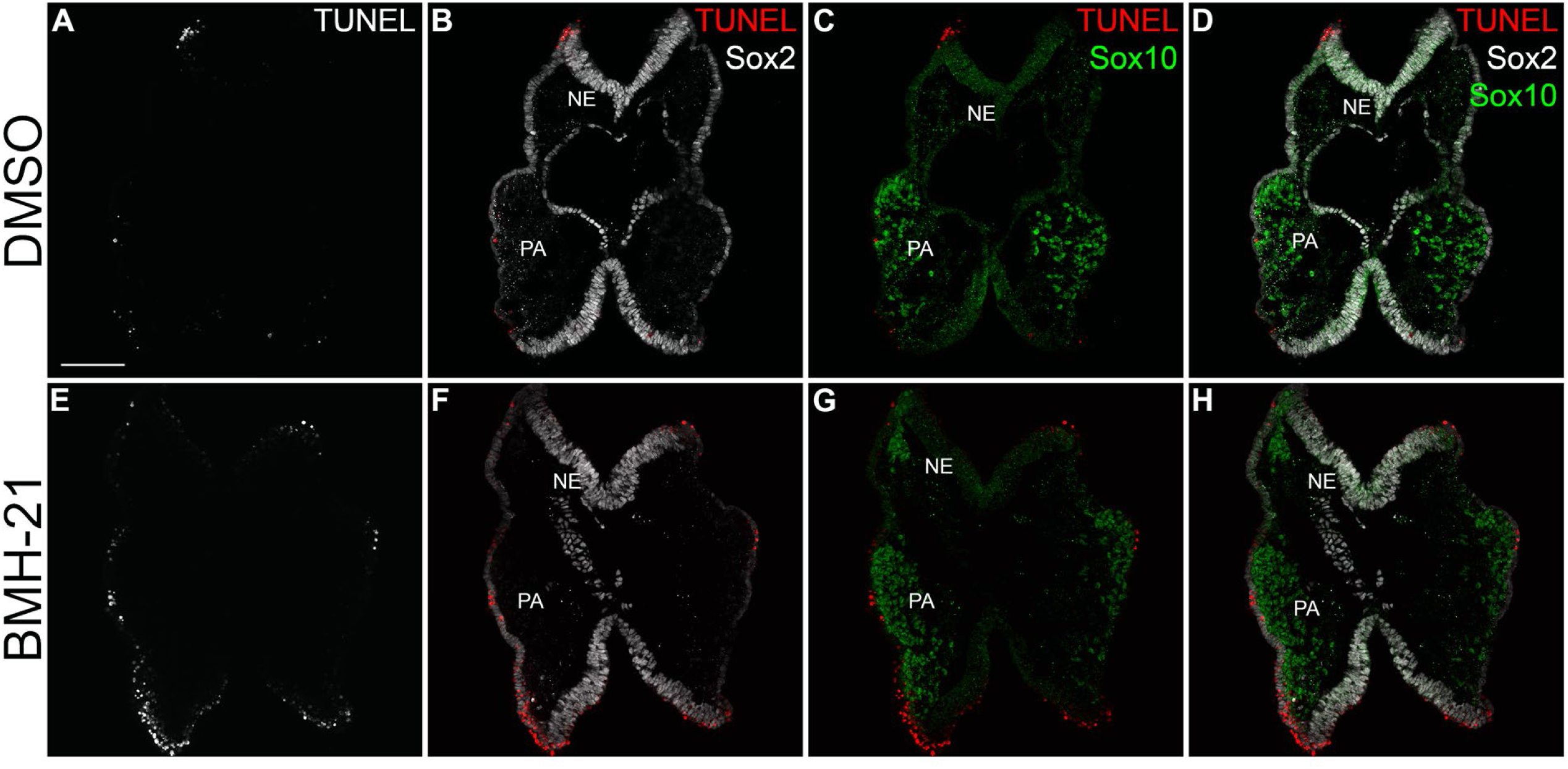
Inhibition of Pol I transcription leads to higher apoptosis in the neuroepithelium and neural crest cells. Transverse sections of DMSO (A-D) and BMH-21(E-F) treated embryos at E8.5 stained for TUNEL (white in A and E, red in B-D and F-H), Sox2 (white in B-D and F-H) and Sox10 (green in C, D, G and H) show that chemical inhibition of Pol I leads to apoptosis, especially in Sox2 positive neuroepithelium and Sox10 positive neural crest cells. Abbreviations: NE, neuroepithelium; PA, pharyngeal arches. Scale bar = 120 µm.

**Fig. S3.**
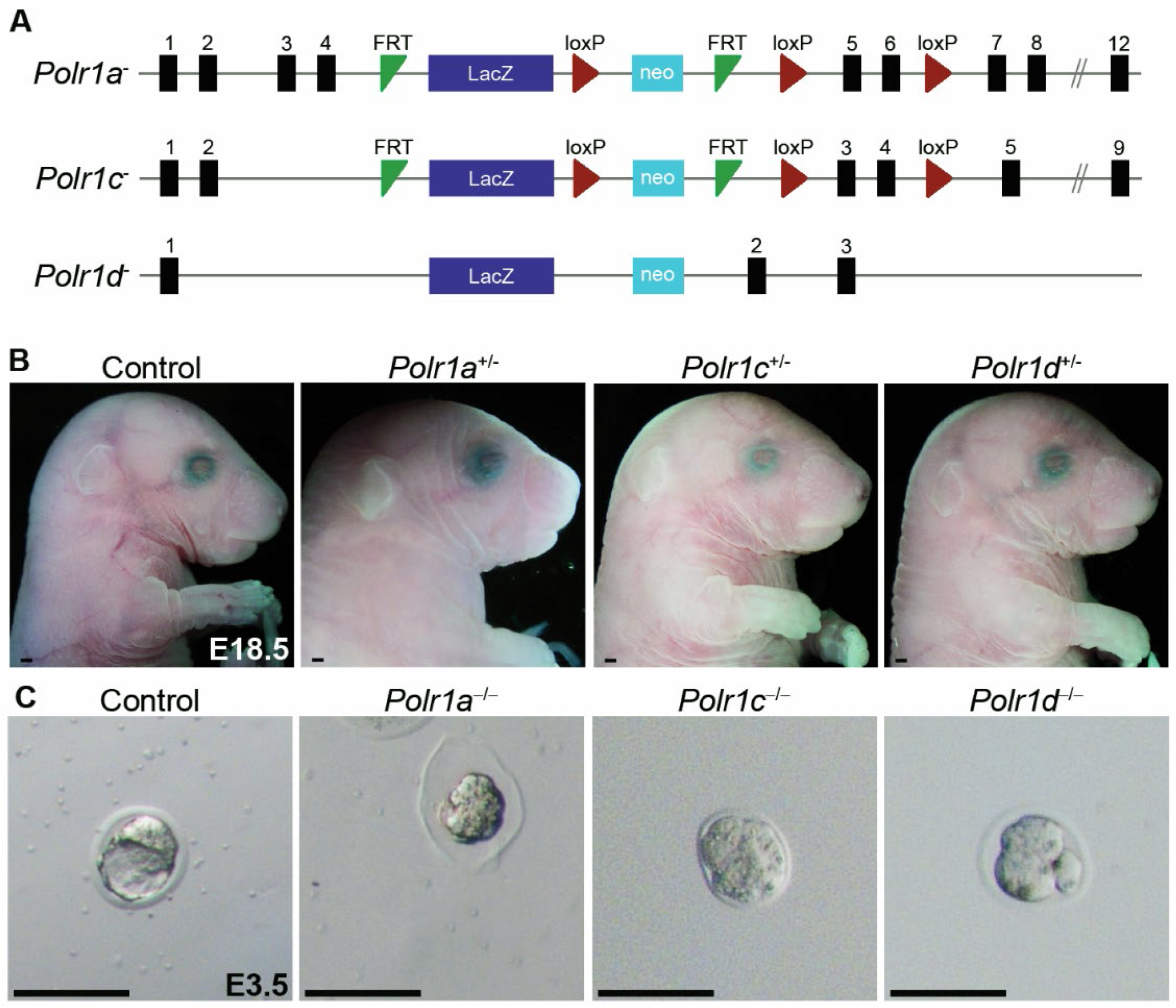
Generation of *Polr1a, Polr1c* and *Polr1d* mutant mice. (A) A LacZ-neo cassette containing loxP sites flanking critical exons of *Polr1a* and *Polr1c* was used to generate *Polr1a^+/-^* and *Polr1c^+/-^* alleles. The *Polr1d^+/-^*allele was generated by disrupting the *Polr1d* gene with a LacZ-neo insert. (B) Heterozygous mutants of *Polr1a*, *Polr1c* and *Polr1d* are indistinguishable from wild-type controls indicating a single copy of *Polr1a*, *Polr1c* and *Polr1d* is sufficient for embryonic development. Scale bar = 500 µm (C) Null mutants of *Polr1a*, *Polr1c* and *Polr1d* survive until the uncompacted morula stage at E2.5 and are fragmented by E3.5 while wildtype embryos proceed to the blastocyst stage indicating *Polr1a*, *Polr1c* and *Polr1d* are required for pre-implantation embryo survival. Scale bar = 100 µm.

**Fig. S4.**
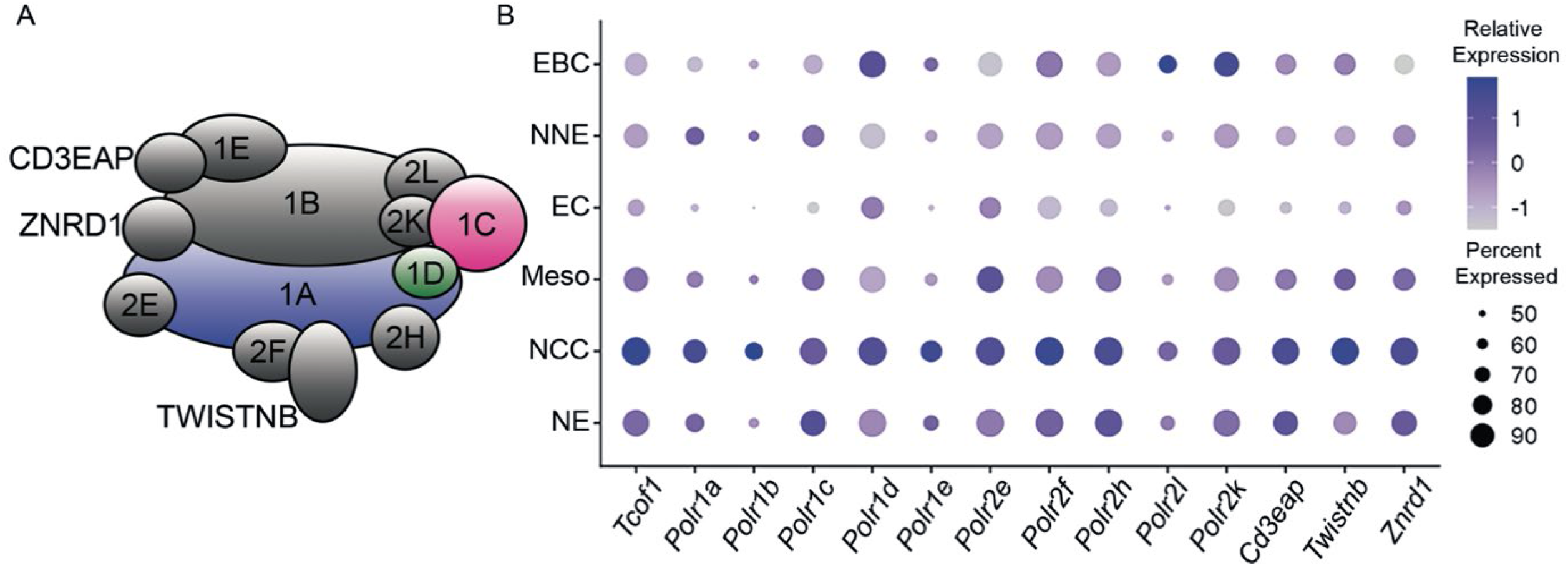
Pol I subunits are expressed highly in NCC. A) Schematic of RNA Polymerase I subunits. B) Single cell RNA-seq analysis identifies *Tcof1* and RNA Polymerase I subunit transcripts to be highly expressed in neuroepithelium and NCC compared to other tissues in the craniofacial region of E8.5 mouse embryos. The size of the circle represents the percent of cells in a population expressing the transcript of interest, while color intensity represents the relative level of transcripts expressed in a cell population (see Methods). Abbreviations: EBC, embryonic blood cells; EC, endothelial cells; Meso, mesoderm; NCC, neural crest cells; NE, neuroepithelium; NNE, non-neural ectoderm.

**Fig. S5.**
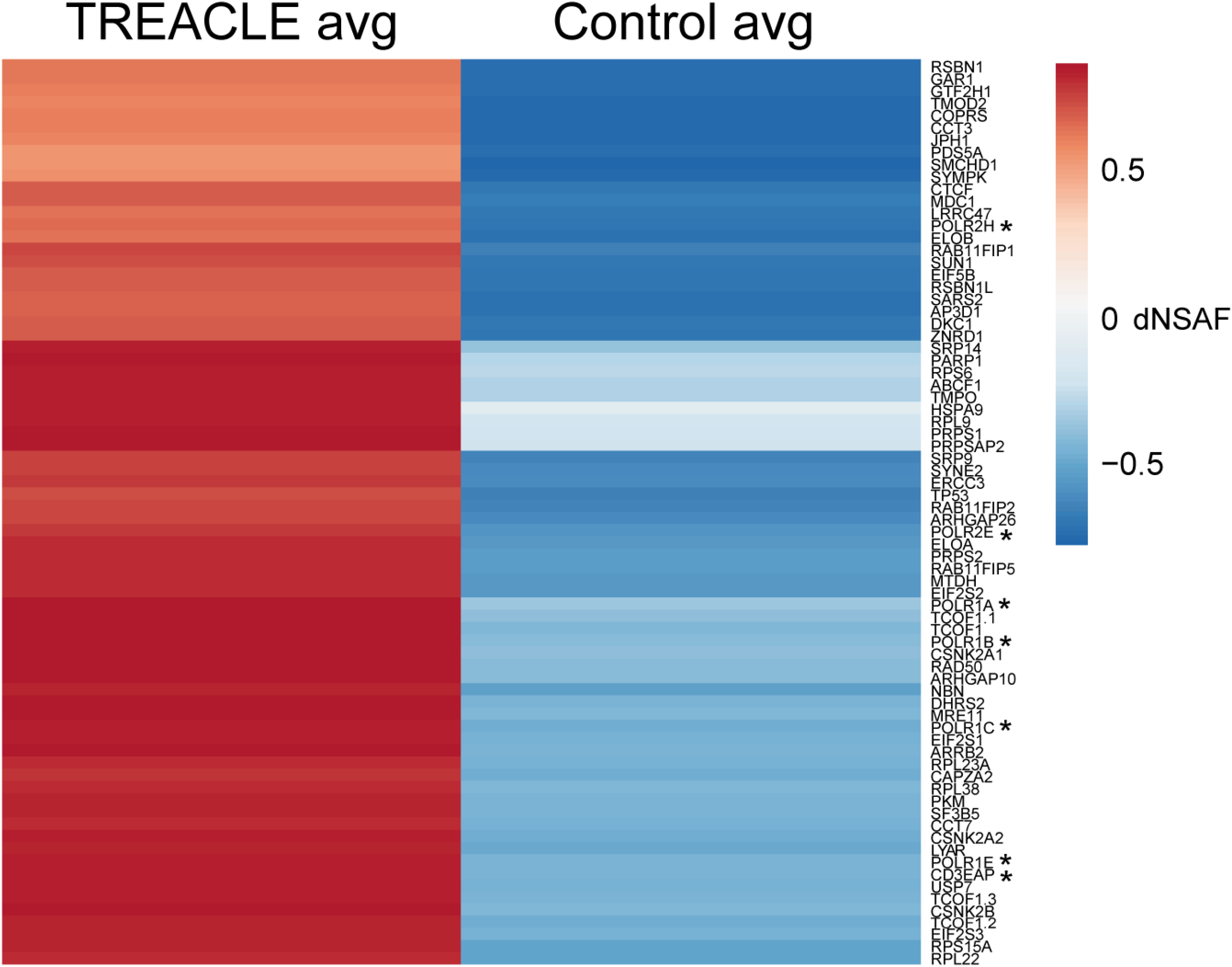
Proteins identified by multidimensional protein identification technology with TREACLE as the bait. Heat map showing the spectral abundance of proteins pulled down with FLAG-tagged TREACLE in both antibody and IgG (control) immunoprecipitation conditions, expressed as distributed normalized spectral abundance factor (dNSAF). This demonstrates the specificity of TREACLE binding to its target proteins. * Pol I protein subunits.

**Fig. S6.**
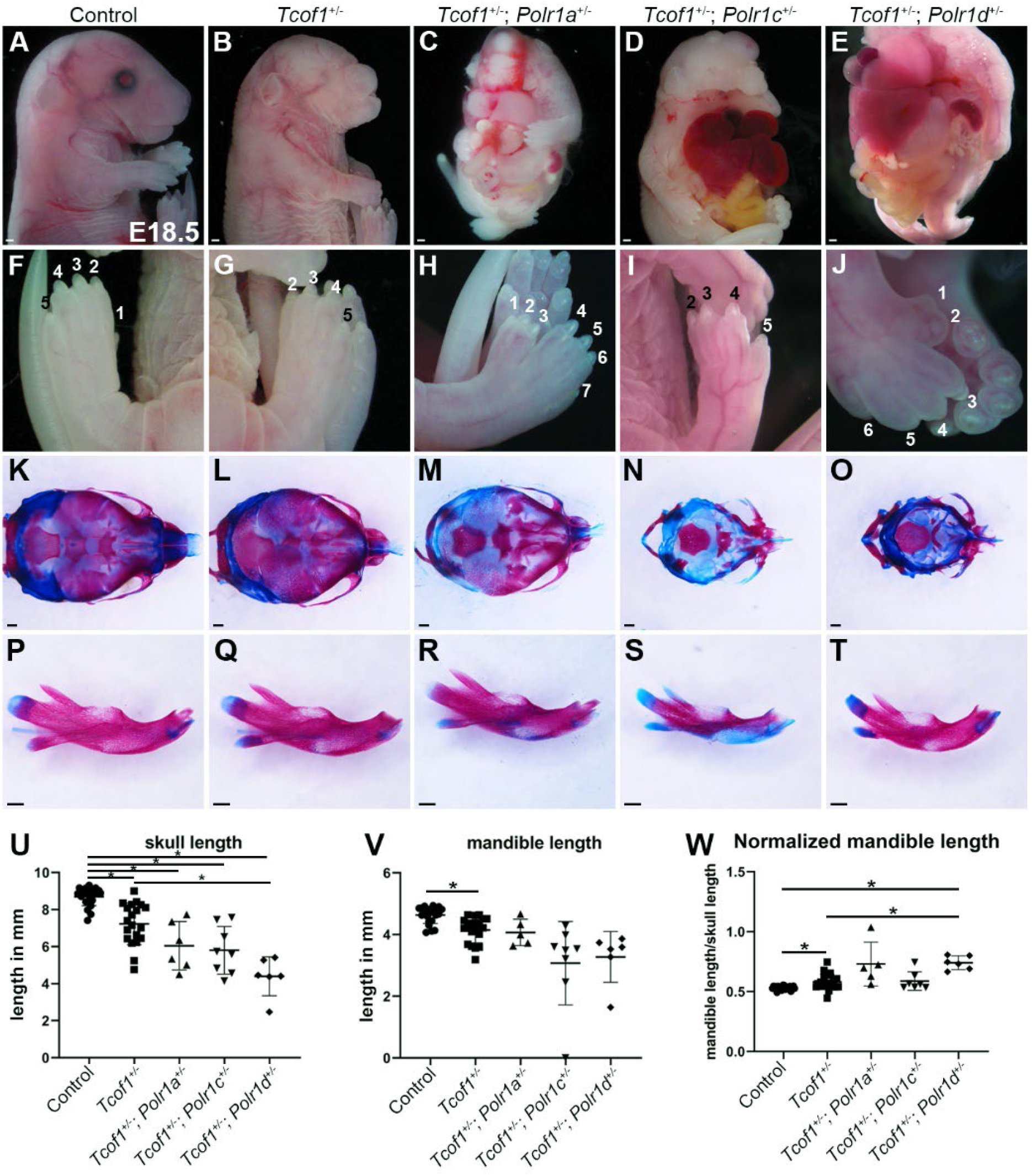
Double mutants of *Tcof1* with *Polr1a*, *Polr1c,* and *Polr1d* exhibit thoracoschisis and digit defects. (A-E) Compared to controls and *Tcof1^+/-^* embryos, *Tcof1^+/-^; Polr1a^+/-^, Tcof1^+/-^; Polr1c^+/-^ and Tcof1^+/-^; Polr1d^+/-^* double mutants exhibit variably penetrant thoracoschisis as evidenced by herniation of lung, liver and gut at variable penetrance. (F-J) *Tcof1^+/-^; Polr1a^+/-^, Tcof1^+/-^; Polr1c^+/-^ and Tcof1^+/-^; Polr1d^+/-^* double mutants exhibit digit defects including duplication of digit 1 (H,J) and shorter, broader digits (I). (K-O) Alcian blue and alizarin red stained skeletons reveal hypoplasia of the skull in *Tcof1^+/-^*, *Tcof1^+/-^; Polr1a^+/-^, Tcof1^+/-^; Polr1c^+/-^* and *Tcof1^+/-^; Polr1d^+/-^* mutants, quantified in (U). (P-T) Dissected mandibles from control, *Tcof1^+/-^*, *Tcof1^+/-^; Polr1a^+/-^, Tcof1^+/-^; Polr1c^+/-^* and *Tcof1^+/-^; Polr1d^+/-^* mutants show reduced mandible length, quantified in (V). (W) Quantification of the size of the mandible relative to the skull demonstrates that the proportion of the mandible relative to skull size is variable in mutant mice. * = p<0.05. Scale bar = 500 µm

**Fig. S7.**
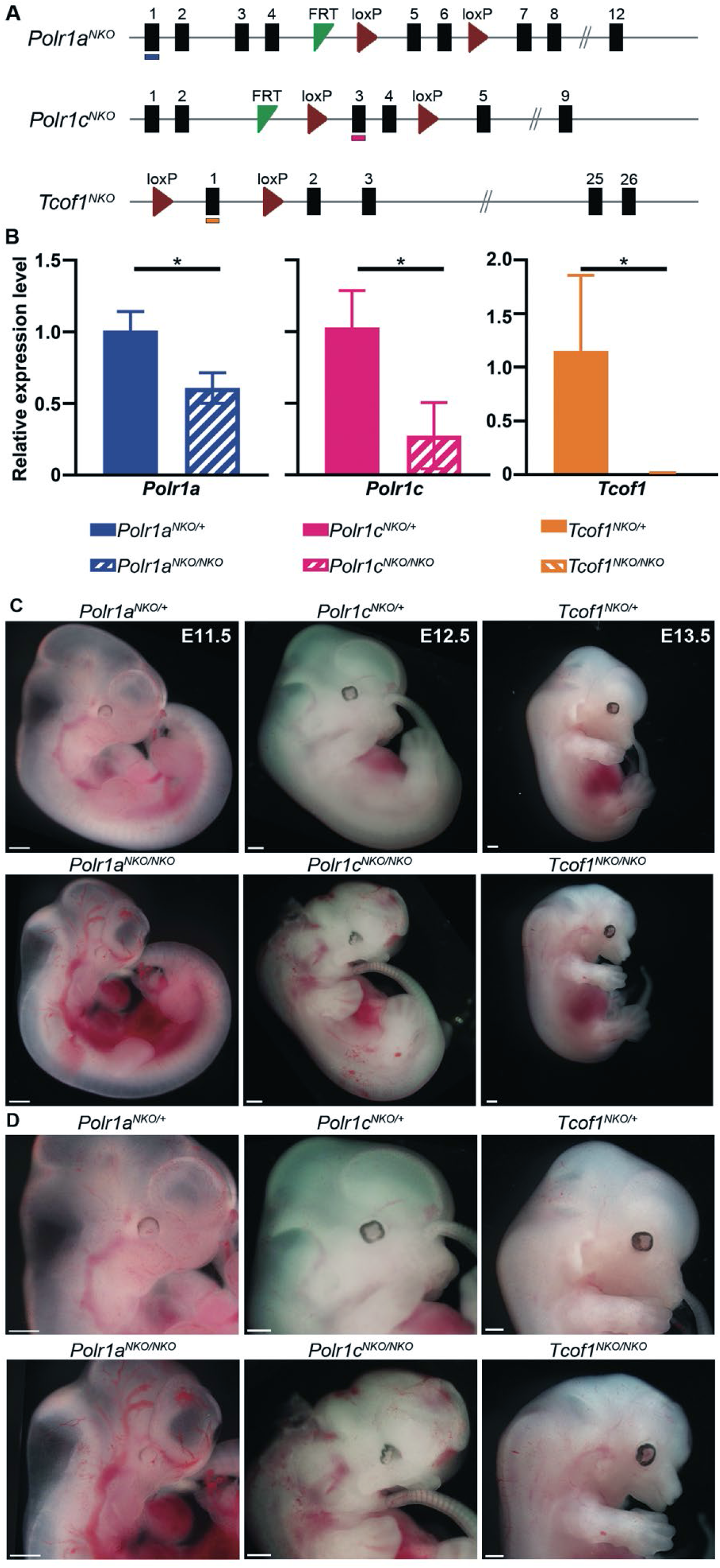
NCC-specific mutants of *Polr1a*, *Polr1c* and *Tcof1* exhibit mid-gestation lethality. (A) NCC-specific knockouts of *Polr1a (Polr1a^NKO^)*, *Polr1c (Polr1c^NKO^)* and *Tcof1 (Tcof1^NKO^)* were generated by flanking critical exons with loxP sites and breeding the floxed allelic mice with *Wnt1-Cre* transgenic mice. Exons examined by qPCR in (B) are underlined in each corresponding construct. (B) qPCR reveals reduced expression of *Polr1a*, *Polr1c,* and *Tcof1* transcripts in sorted NCC from *Polr1a^NKO/NKO^*, *Polr1c^NKO/NKO^*, and *Tcof1^NKO/NKO^* embryos, respectively, compared to their control littermates at E9.5. *indicates p<0.05, Student’s t-test. (C) While a single copy of *Polr1a*, *Polr1c* and *Tcof1* in NCC is sufficient for embryonic development, knocking out both copies of *Polr1a*, *Polr1c* and *Tcof1* from NCC results in midgestation lethality. (D) Higher magnification images of (C). Scale bar = 500 µm.

**Figure S8.**
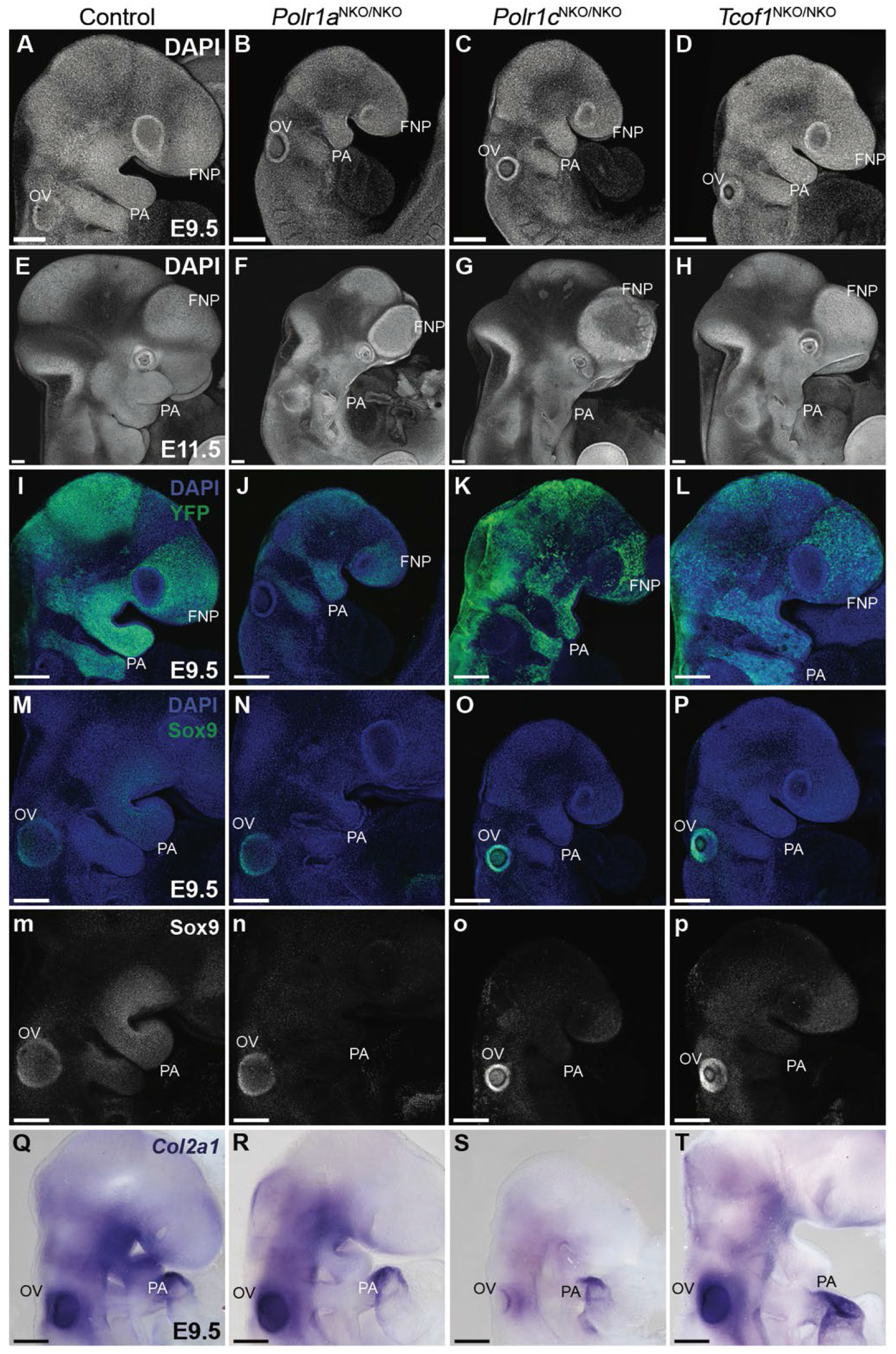
Craniofacial defects in *Polr1a^NKO/NKO^*, *Polr1c^NKO/NKO^* and *Tcof1^NKO/NKO^* mice. (A-H) DAPI staining of control, *Polr1a^NKO/NKO^*, *Polr1c^NKO/NKO^* and *Tcof1^NKO/NKO^* embryos at E9.5 and E11.5 shows hypoplastic pharyngeal arches and frontonasal prominences in *Polr1a^NKO/NKO^*, *Polr1c^NKO/NKO^* and *Tcof1^NKO/NKO^* embryos. (I-L) Analysis of the NCC lineage with RosaeYFP indicates that fewer NCC migrate to the pharyngeal arches in *Polr1a^NKO/NKO^*, *Polr1c^NKO/NKO^* and *Tcof1^NKO/NKO^* at E9.5. (M-P) Immunostaining for Sox9, a marker indicative of NCC migration and differentiation to chondrocytes (green in M-P and gray in m-p). Sox9 is significantly reduced in the pharyngeal arches of *Polr1a^NKO/NKO^*, *Polr1c^NKO/NKO^* and *Tcof1^NKO/NKO^* embryos, indicating a reduction in the number of NCC precursors necessary for craniofacial cartilage development. (Q-T) Consistent with this, Type II collagen *Col2a1* transcription is drastically reduced in the pharyngeal arches of *Polr1a^NKO/NKO^*, *Polr1c^NKO/NKO^* and *Tcof1^NKO/NKO^* embryos at E9.5. Abbreviations. FNP, Frontonasal prominence; NE, neuroepithelium; OV, Otic vesicle. PA, pharyngeal arches; Scale bar = 200 µm.

**Figure S9.**
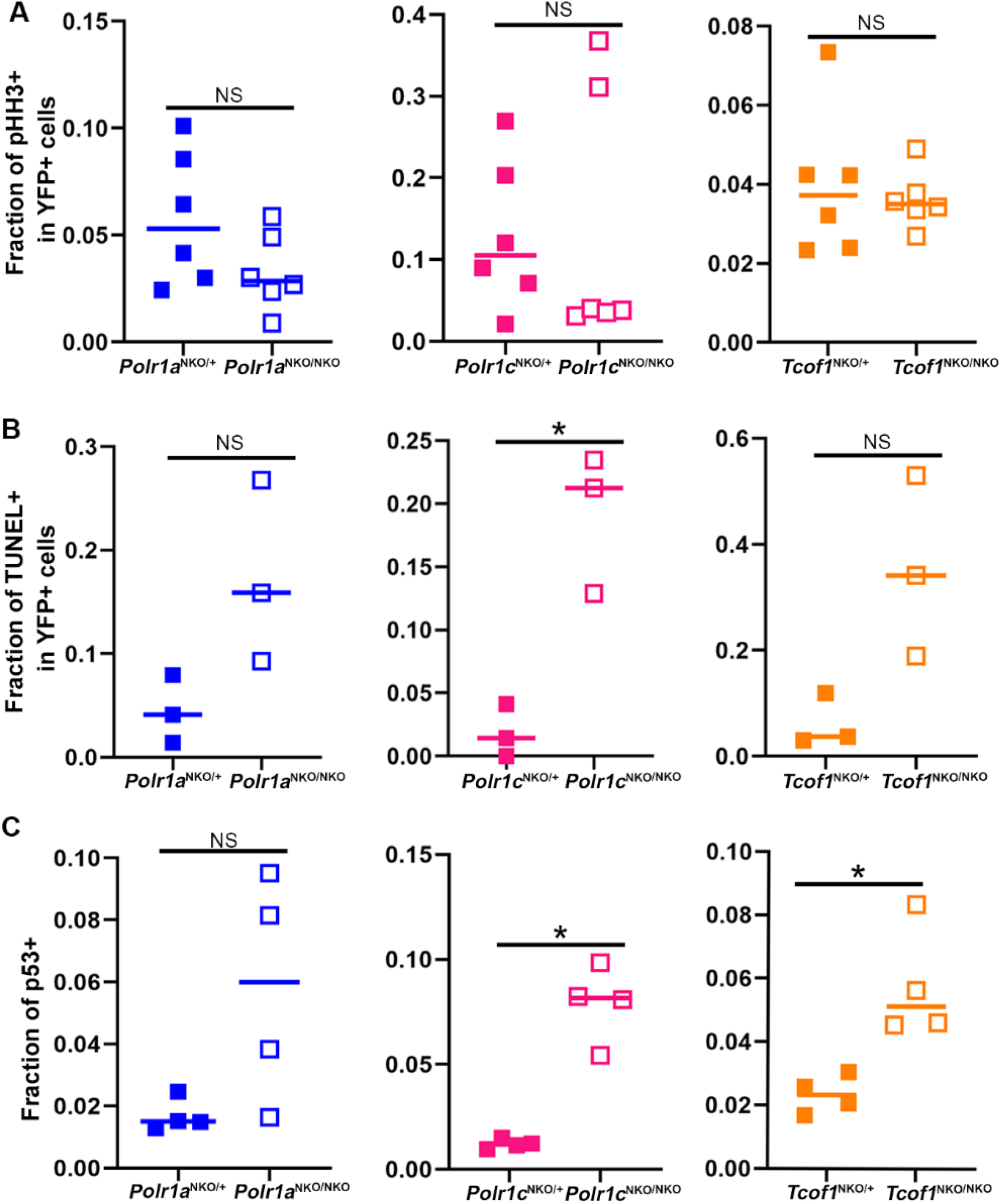
Quantification of pHH3, TUNEL, and p53 in NCC of *Polr1a^NKO/NKO^*, *Polr1c^NKO/NKO^* and *Tcof1^NKO/NKO^* embryos compared to *Polr1a^NKO/+^*, *Polr1c^NKO/+^* and *Tcof1^NKO/+^* controls. (A) Quantification of pHH3+; YFP+ cells demonstrates that pHH3 staining in the NCC at E9.5 tends to be less in *NKO/NKO* mutants relative to littermate controls, however this difference was not statistically significant. (B) Quantification of TUNEL+;YFP+ cells reveals increased cell death in *NKO/NKO* mutants. This trend was statistically significant in *Polr1c^NKO/NKO^* embryos. (C) Similarly, quantification of p53+ cells revealed increased levels of p53 in *NKO/NKO* mutants. This trend was statistically significant in *Polr1c^NKO/NKO^* and *Tcof1^NKO/NKO^* embryos relative to their littermate controls. * indicates p<0.05, Student’s t-test.

**Figure S10.**
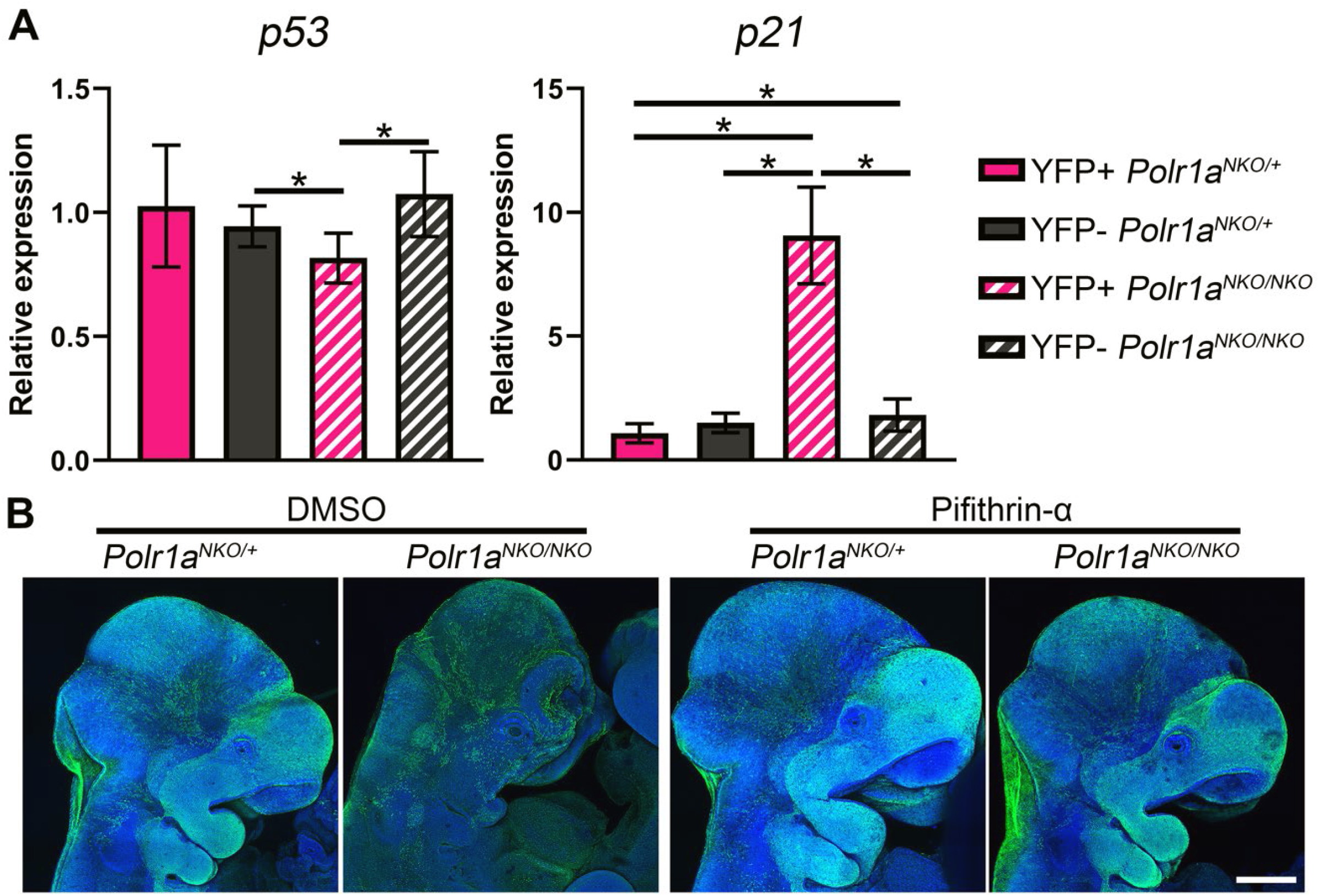
Quantification of *p53* and *p21* transcript in NCC and non-NCC of *Polr1a^NKO/NKO^* embryos compared to *Polr1a^NKO/+^* controls. (A) qPCR for *p53* demonstrates that *p53* levels are not significantly changed in NCC versus non-NCC in control embryos (YFP+ *Polr1a^NKO/+^* vs. YFP-*Polr1a^NKO/+^*) while *p53* transcript levels are significantly downregulated in NCC versus non-NCC in mutant embryos (YFP+ *Polr1a^NKO/NKO^* vs. YFP-*Polr1a^NKO/NKO^*) and versus non-NCC in controls (YFP+ *Polr1a^NKO/NKO^* vs. YFP-*Polr1a^NKO/+^*). However, *p53* levels in the NCC population between YFP+ *Polr1a^NKO/NKO^* mutants and *YFP+ Polr1a^NKO/+^* controls was not significantly changed. qPCR for *p21* demonstrates that *p21* is significantly upregulated in YFP+ *Polr1a^NKO/NKO^* mutants compared to all other cell populations examined. *indicates p<0.05, Welch’s ANOVA, Dunnett’s T3 multiple comparisons test. (B) Immunostaining for RosaeYFP to assess NCC lineage in DMSO and pifithrin-a treated *Polr1a^NKO/NKO^* and control embryos indicates that pifithrin-a treatment rescues the pharyngeal arch volumes as well as NCC population in *Polr1a^NKO/NKO^* embryos. Scale bar = 400 µm.

**Figure S11.**
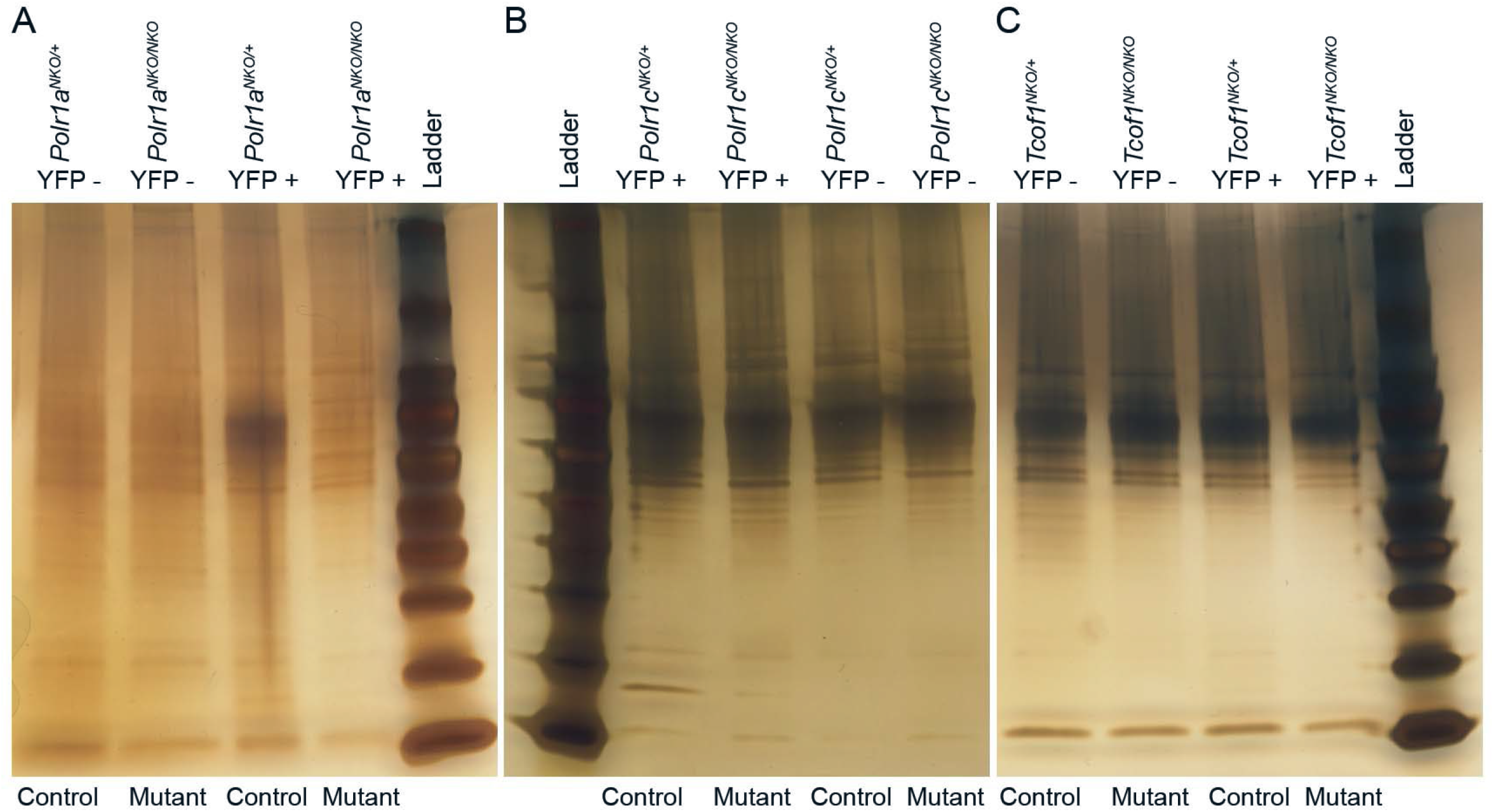
Total protein is significantly reduced in the NCC of *Polr1a^NKO/NKO^*, *Polr1c^NKO/NKO^* and *Tcof1^NKO/NKO^* mice. (A-C) Silver staining reveals that total protein levels in equal numbers of NCC (YFP+) is slightly higher compared to non-NCC (YFP-) in E10.5 control embryos as observed by the number of bands and intensity of bands in silver-stained gels. However, protein levels in NCC (YFP+) are comparable to non-NCC (YFP-) in mutant *Polr1a^NKO/NKO^*, *Polr1c^NKO/NKO^* and *Tcof1^NKO/NKO^* embryos. Compared to control NCC (YFP+), protein expression is significantly lower in mutant *Polr1a^NKO/NKO^*, *Polr1c^NKO/NKO^* and *Tcof1^NKO/NKO^* NCC (YFP+).

**Figure S12.**
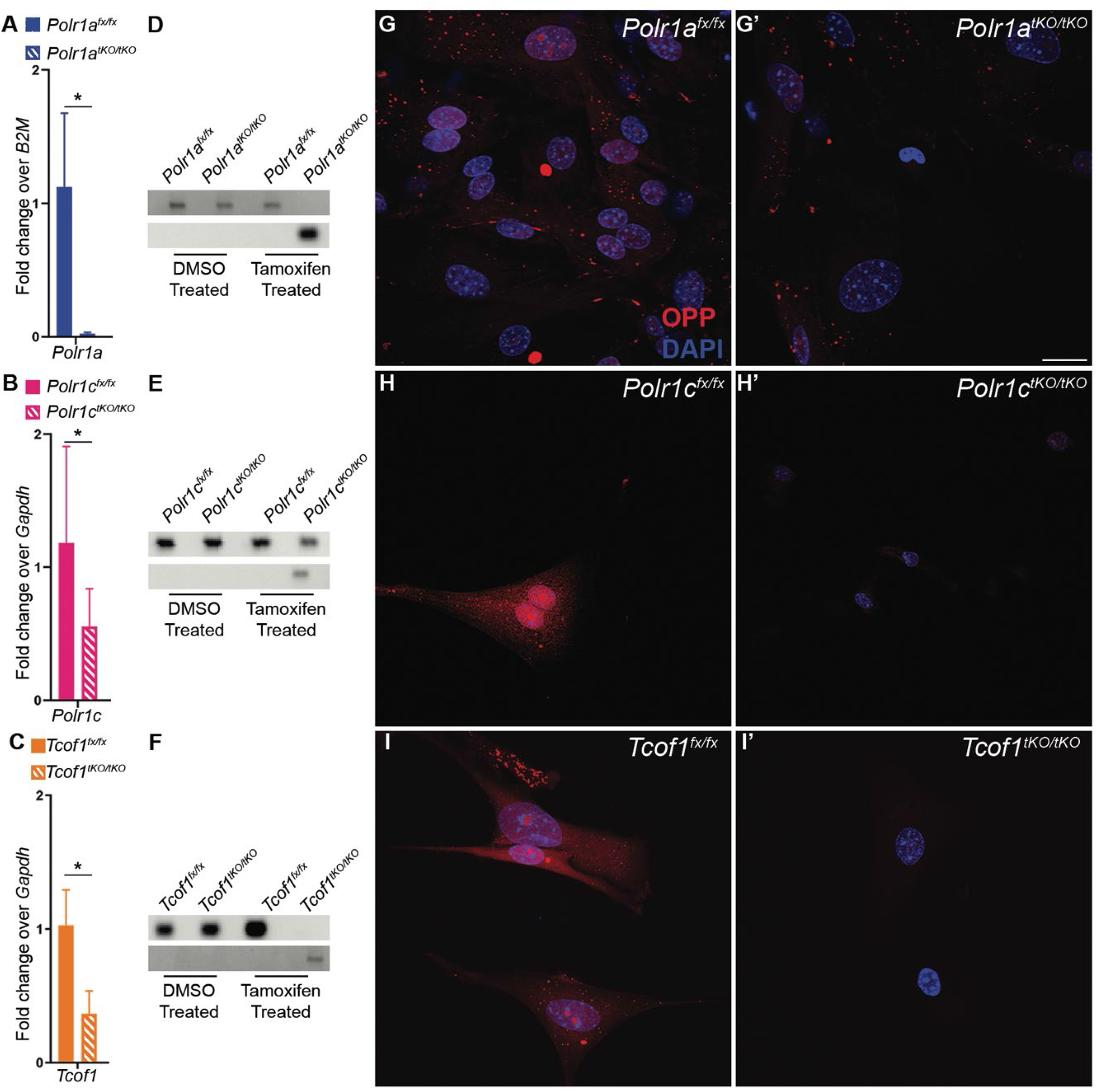
*Polr1a^tKO/tKO^*, *Polr1c^tKO/tKO^* and *Tcof1^tKO/tKO^* mouse embryonic fibroblast cells have defects in protein synthesis. (A-C) Tamoxifen-treated MEFs generated from *Polr1a^tKO/tKO^*, *Polr1c^tKO/tKO^* and *Tcof1^tKO/tKO^* mutant embryos have reduced expression of *Polr1a*, *Polr1c* and *Tcof1* transcripts, respectively. * indicates p<0.05, Student’s t-test. (D-F) Recombined DNA is specifically present in tamoxifen treated *Polr1a^tKO/tKO^*, *Polr1c^tKO/tKO^* and *Tcof1^tKO/tKO^* MEFs. (H-I’) New protein synthesis is significantly reduced in *Polr1a^tKO/tKO^*, *Polr1c^tKO/tKO^* and *Tcof1^tKO/tKO^* as observed by OPP staining. Scale bar = 140 µm.

**Figure S13.**
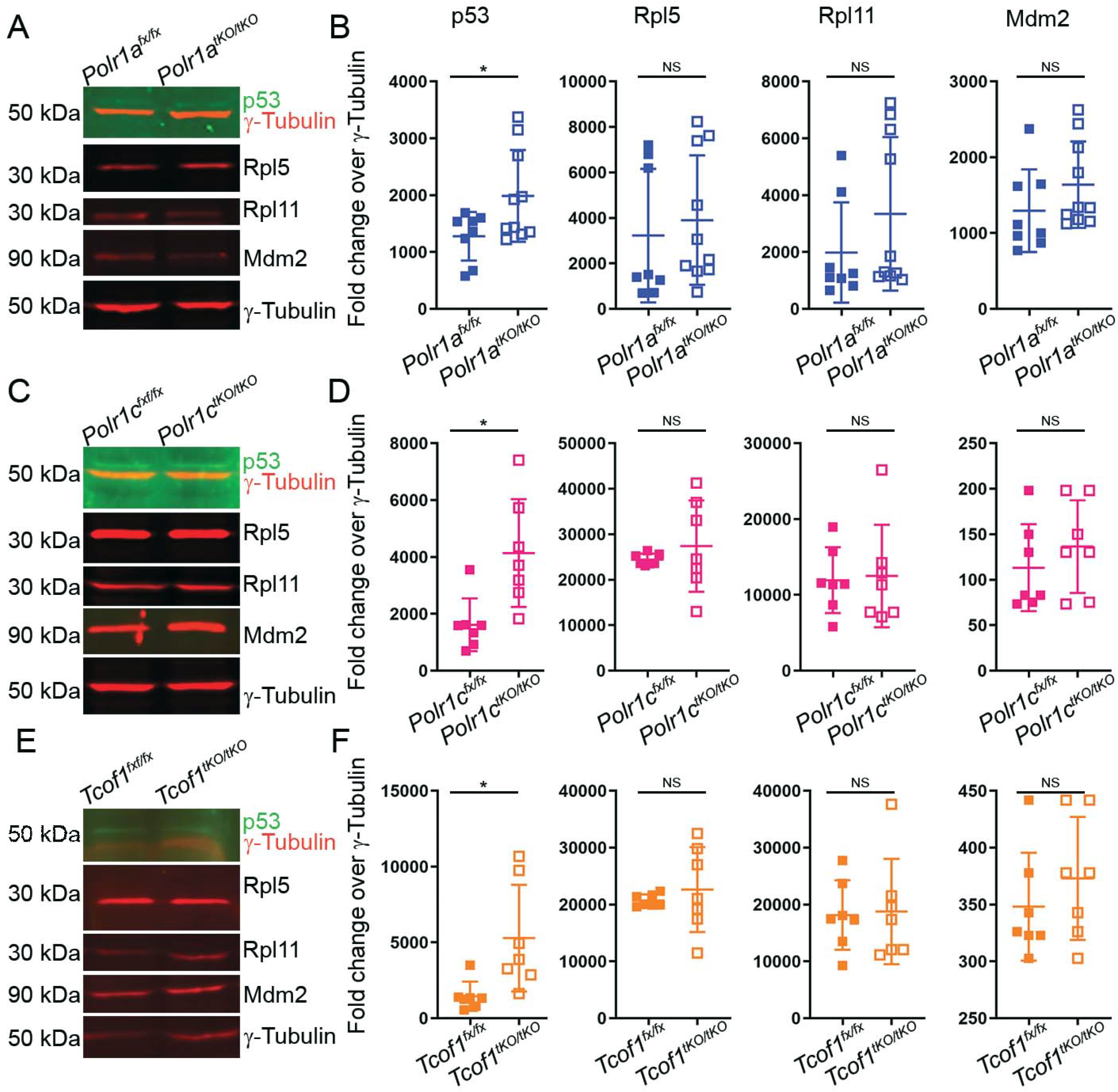
p53 upregulation is observed in *Polr1a^tKO/tKO^*, *Polr1c^tKO/tKO^* and *Tcof1^tKO/tKO^* mouse embryonic fibroblast cells. (A-F) *Polr1a^tKO/tKO^*, *Polr1c^tKO/tKO^* and *Tcof1^tKO/tKO^* MEFs have increased p53 expression/accumulation. In contrast, Rpl5, Rpl11 and Mdm2 levels are not significantly changed between controls and *Polr1a^tKO/tKO^*, *Polr1c^tKO/tKO^* and *Tcof1^tKO/tKO^* MEFs. ɣ-Tubulin was used as a housekeeping gene and fold change was calculated as a ratio of band intensities. * indicates p<0.05, Student’s t-test. Abbreviations: NS, not significant.

**Figure S14.**
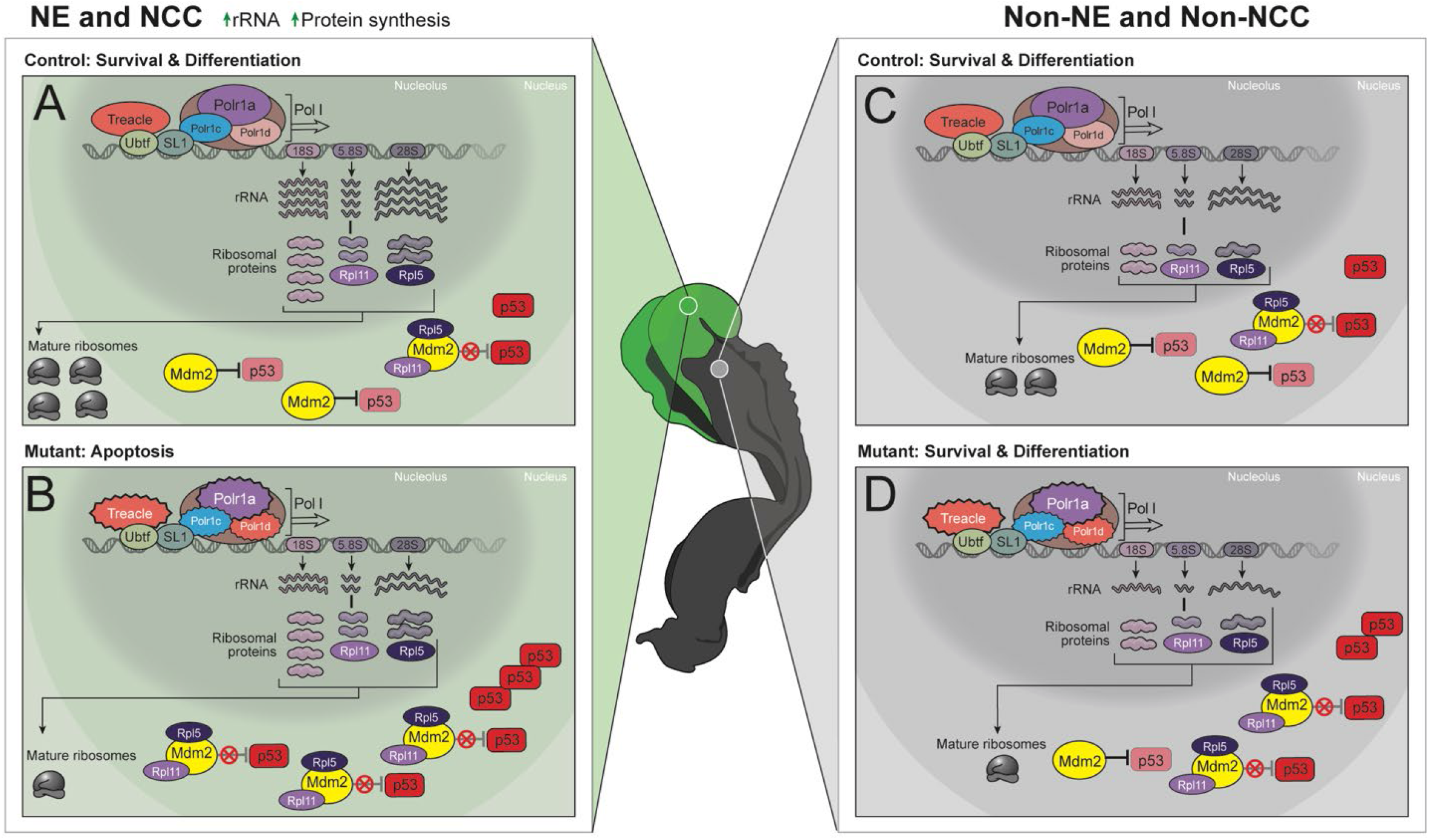
NCC are more sensitive to disruptions in rRNA transcription, which leads to increased susceptibility to p53-dependent cell death. (A) rRNA and ribosomal proteins are maintained in balanced quantities for proper ribosome assembly. The neuroepithelium (NE) and NCC are highly proliferative and have elevated levels of rDNA transcription relative to surrounding tissues, and thus high levels of ribosomal proteins. During normal cell growth and proliferation, Mdm2 protein binds to, and ubiquitinates p53, targeting it for degradation. This typically keeps p53 at low levels and maintains cell survival. (B) Increased levels of rDNA transcription make NE and NCC highly susceptible to disruptions in Pol I mediated transcription. Disruption of rRNA synthesis due to deletion in Pol I subunits Polr1a, Polr1c, Polr1d, or associated factor Treacle, results in increased free ribosomal proteins and thus binding of Rpl5 and Rpl11 to Mdm2. This inhibits Mdm2 binding and degradation of p53 resulting in p53 protein accumulation. Increased p53 results in NCC apoptosis and craniofacial malformations. (C) Cells with lower proliferative capacity than NE and NCC, have lower rDNA transcription, less rRNA and ribosomal proteins. (D) Upon disruption of rRNA synthesis in these cells, the levels of free ribosomal proteins remain low with little binding of Rpl5 and Rpl11 to Mdm2. Thus, Mdm2 continues to bind to and ubiquitinate p53 targeting it for degradation, which helps to keep p53 protein levels low in support of cell survival.

**Table S1.**
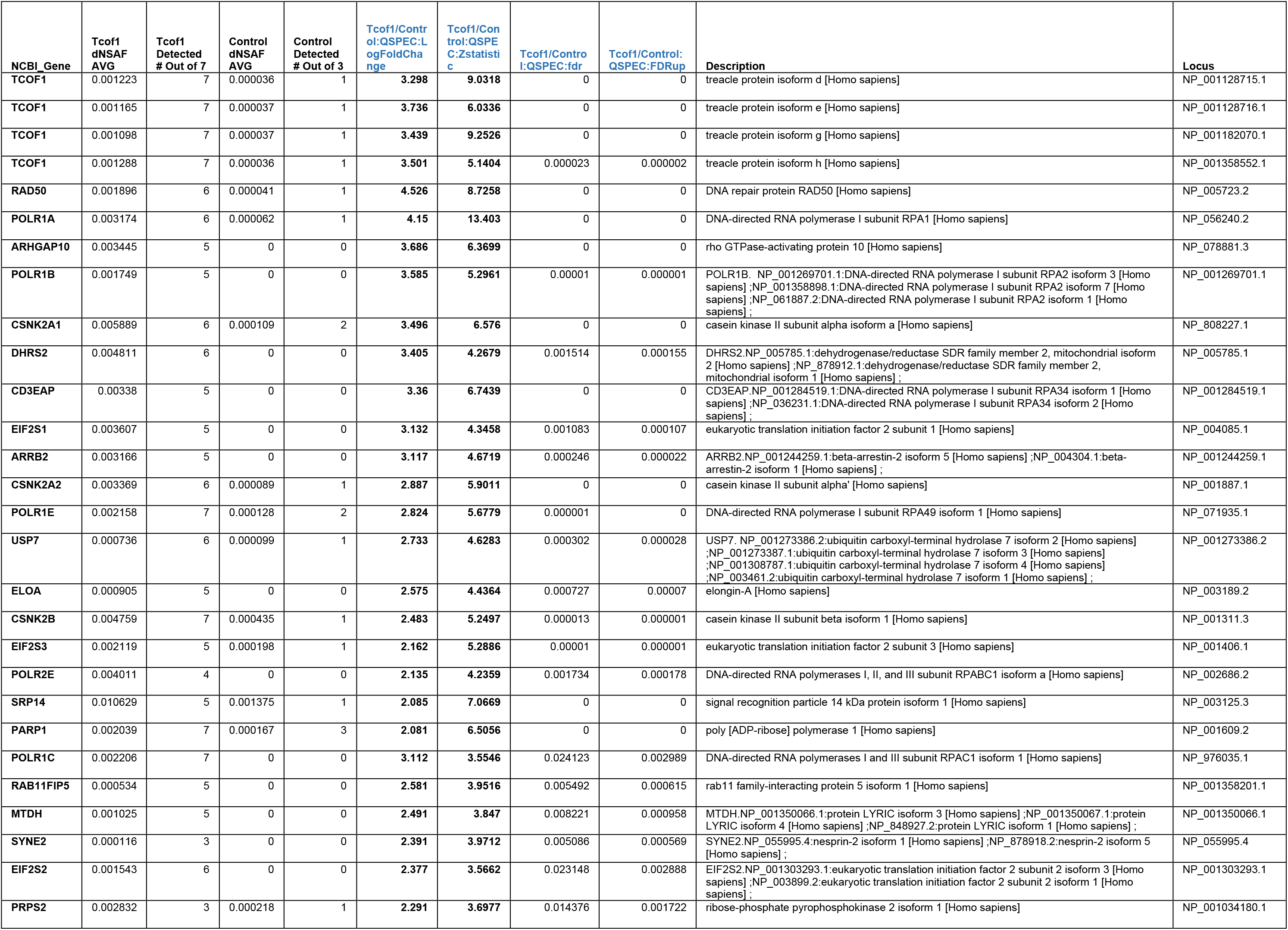
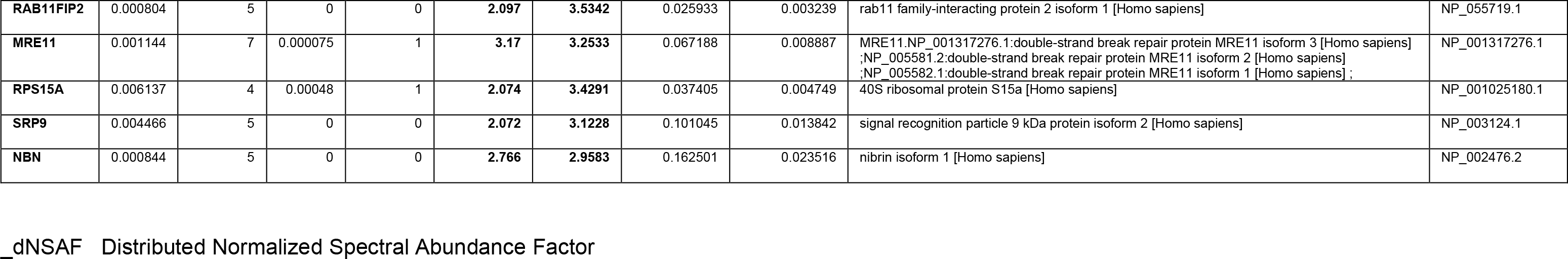
Top 30 proteins identified by multidimensional protein identification technology with TCOF1/TREACLE as the bait or the 293 Control.

**Table S2.**
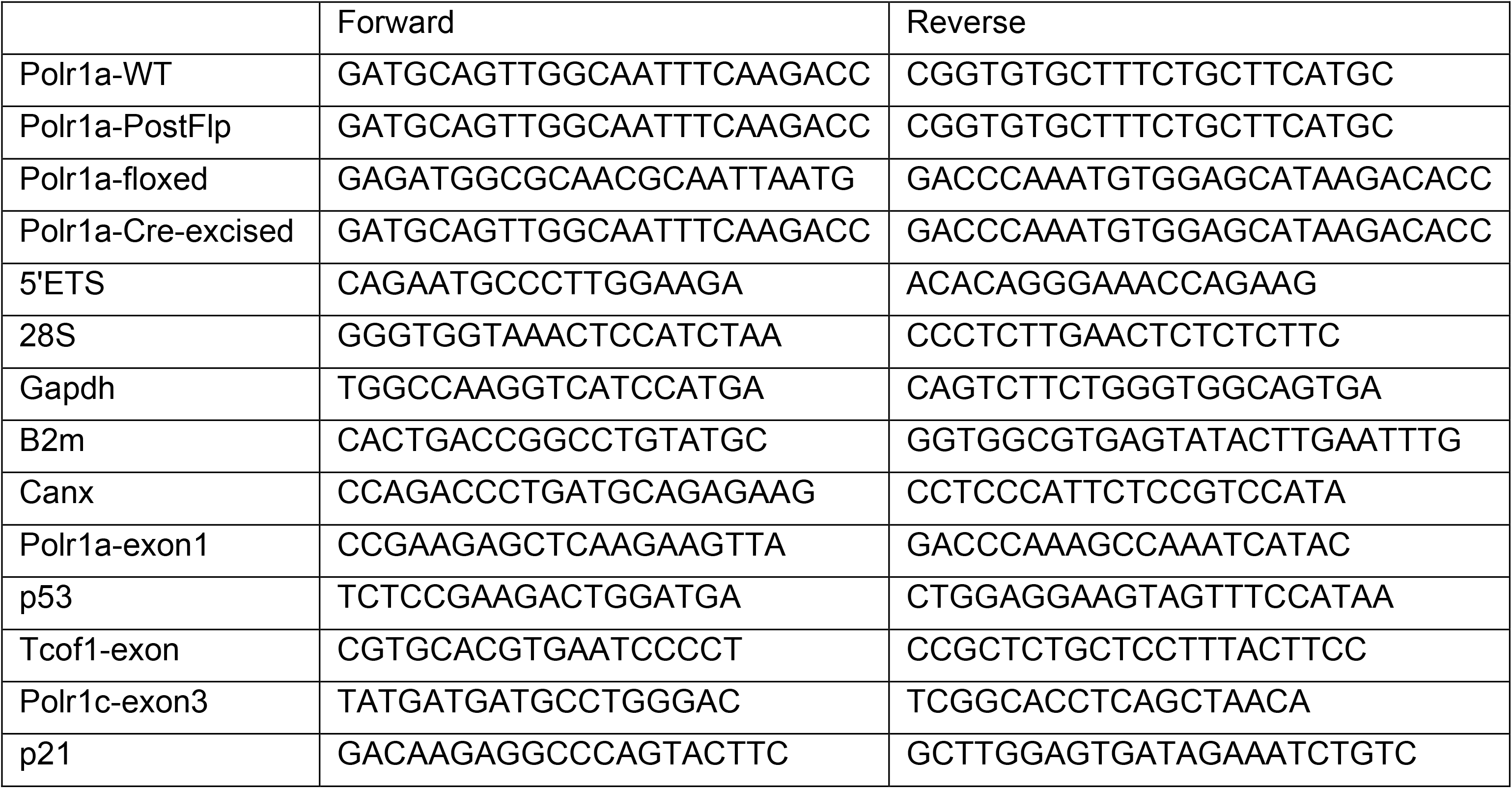
List of primers used for genotyping and RT-qPCR.

